# Dietary protein source alters gut physiology through direct and microbiota-mediated effects

**DOI:** 10.1101/2025.01.09.632171

**Authors:** Ayesha Awan, J. Alfredo Blakeley-Ruiz, Clara Griggs, Shrey D. Thaker, Hope Chronis, Alexandria Bartlett, Lorraine Danowski, Josephine Connolly-Schoonen, David C. Montrose, Manuel Kleiner

## Abstract

Dietary protein source differentially impacts long-term health outcomes, yet the specific host responses underlying these health effects remain unclear, including which are direct effects of protein source and which are microbiota-mediated. Using metagenome-informed metaproteomics, we determined that protein source altered the abundance of host proteins related to proteolysis, immunity, and the gut barrier as well as microbial function in mice. These stool host responses further reflected site-specific responses throughout the intestinal tract. Additionally, while some host responses were microbiota-dependent, others were microbiota-independent. These effects of dietary protein source on the host and microbiota translated to humans. Notably, microbial enzymes related to mucin degradation increased consistently in response to egg white protein in both mice and humans. Our results show that protein source impacts multiple aspects of host physiology through microbiota-dependent and independent routes, providing essential insights into mechanisms underpinning protein source-driven differences in health outcomes.

## Introduction

Despite the importance of protein as a major macronutrient, and the recent change in dietary habits toward higher intake of protein and purified protein supplements^1^, research on the functional impacts of dietary protein from different sources has lagged behind that for other dietary components such as fat and fiber^2^. This is a critical gap, as dietary proteins from different sources differentially affect long-term health outcomes in large human populations^3–7^. For example, increased mortality associated with dietary proteins from specific animal sources, including red meat and eggs, is significantly improved by substituting animal protein with plant protein^4,5,8^. It has been suggested that these impacts on health are mediated by differences in the digestibility of dietary protein from different sources and the subsequent interactions between undigested protein residues and the gut microbiota^2,9^. Previous studies comparing casein and soy protein found that casein led to reduced gut microbial diversity and worsened inflammation in a mouse model of colitis^10^. Compared to casein, soy protein was associated with lower levels of secretory immunoglobulin A, IL-4, IL-13, mucin 1, and mucin 2^11^.

Additionally, studies have shown that specific characteristics of dietary proteins such as the glycosylations associated with egg white and yeast can be metabolized by specific *Bacteroides* species in the gut. Still, the impact of these diet-microbiota interactions on the host remains unknown^12,13^. Research from our lab and others has shown that purified dietary proteins from different sources, including egg whites, brown rice and yeast, are differentially degraded by the host and the gut microbiota, which strongly impacts gut microbial composition and metabolism^13–16^. These effects of dietary protein source on the fate of dietary protein and the gut microbiota do not follow a simple animal versus plant dichotomy and are instead specific to the unique source of protein. However, the impact of these dynamic interactions between dietary protein source, the host, and the gut microbiota on specific aspects of host physiology driving these differential health outcomes is not well characterized. The direct effects of dietary protein source on host physiology, independent of gut microbiota interactions, also remain understudied.

Current approaches used to measure host response to diet-microbiota interactions, including ELISAs and host stool transcriptomics, face limitations such as only measuring a single protein or relying on host mRNA, which is unstable and difficult to enrich from stool^17–20^. Non-invasive stool metaproteomics, with its ability to quantify thousands of host, microbial and dietary proteins in an untargeted manner, is a powerful approach to comprehensively characterize multifaceted changes in gut physiology in the context of host-diet-microbiota interactions^21–25^. The host releases proteins into the mucus layer and the intestinal lumen, which reflect the host’s interactions with the gut microbiota and diet. Quantifying these proteins offers an opportunity to directly characterize the host’s response to diet and microbial perturbation in the gut. In particular, stool samples contain hundreds of host proteins with various functions, including digestive enzymes, structural proteins, immunoglobulins, antimicrobial proteins, signaling proteins, and epithelial proteins, including cell adhesion and tight junction proteins^20,21,23^. Characterizing changes in these host proteins in stool therefore provides an opportunity to determine changes in several aspects of host physiology in a single measurement^21^. However, studies linking host responses in stool samples to site-specific intestinal tissue responses are lacking. Determining concordance between the magnitude and directionality of change of specific host proteins in stool and tissues across different parts of the intestinal tract could help extract the biological signal for host responses in stool, enabling researchers and clinicians to use these stool host proteins to draw meaningful conclusions about gut physiology.

In this study, we determined the impact of dietary protein source on the host using both data-independent acquisition (DIA) and data-dependent acquisition (DDA) metaproteomic analysis of stool and intestinal tissue samples from the distal ileum, cecum, and colon of mice across 2 experiments. In Experiment 1, we characterized tissue and stool responses to dietary protein source and determined how well site-specific gut responses were reflected in the stool host proteome. In Experiment 2, we then distinguished direct dietary protein source-driven effects on the host from effects mediated by diet-microbiota-host interactions by comparing host responses in the presence and absence of the gut microbiota. We further determined whether the effects of dietary protein source in mice translated to humans by comparing the host response in people fed either egg white or pea protein. Our results show that dietary protein source impacts multiple aspects of host physiology in both mice and humans. In mice, we found that these effects span the length of the intestinal tract and that site-specific changes in the ileum, cecum, and colon are captured in the stool host proteome. While some of these effects on the host are dependent on the gut microbiota, others are microbiota-independent and directly related to the dietary protein source. Additionally, we found consistent effects of dietary protein source on the gut microbiota function in both mice and humans. Our results provide insights into the potential mechanistic underpinnings of long-term health consequences of different dietary protein sources, indicating that the physiological consequences of not only protein amount but also source need to be considered when devising health-promoting nutrition strategies.

## Results

### Stool proteome changes reflect host responses along the intestinal tract to dietary protein source

To determine the impact of dietary protein source on host intestinal physiology, we quantified and characterized changes in the host proteome of the ileum, cecum, colon, and stool samples of mice with a conventional microbiota (Fig. 1, Supp. Table 1). We fed the mice defined diets containing a single source of protein from either soy, pea, brown rice, egg whites, or yeast for seven days. These dietary protein sources were selected based on their relevance to animal and human diets^13,26–34^. Across all tissue and stool samples analyzed, we detected 13,000 mouse proteins in at least one sample. Intestinal tissue samples, on average, displayed a higher number of proteins detected in at least one sample (10,000 to 12,500) than stool samples (∼3,000). Using a PERMANOVA on Euclidean distances we found that dietary protein source significantly altered the host proteome across the gut, including the distal ileum (R^2^=0.337, p=0.001), cecum (R^2^=0.413, p=0.001), and colon (R^2^=0.331, p=0.001), as well as in the stool at day 7 (R^2^=0.306, p=0.001) (Fig. 1B).

**Figure 1.**
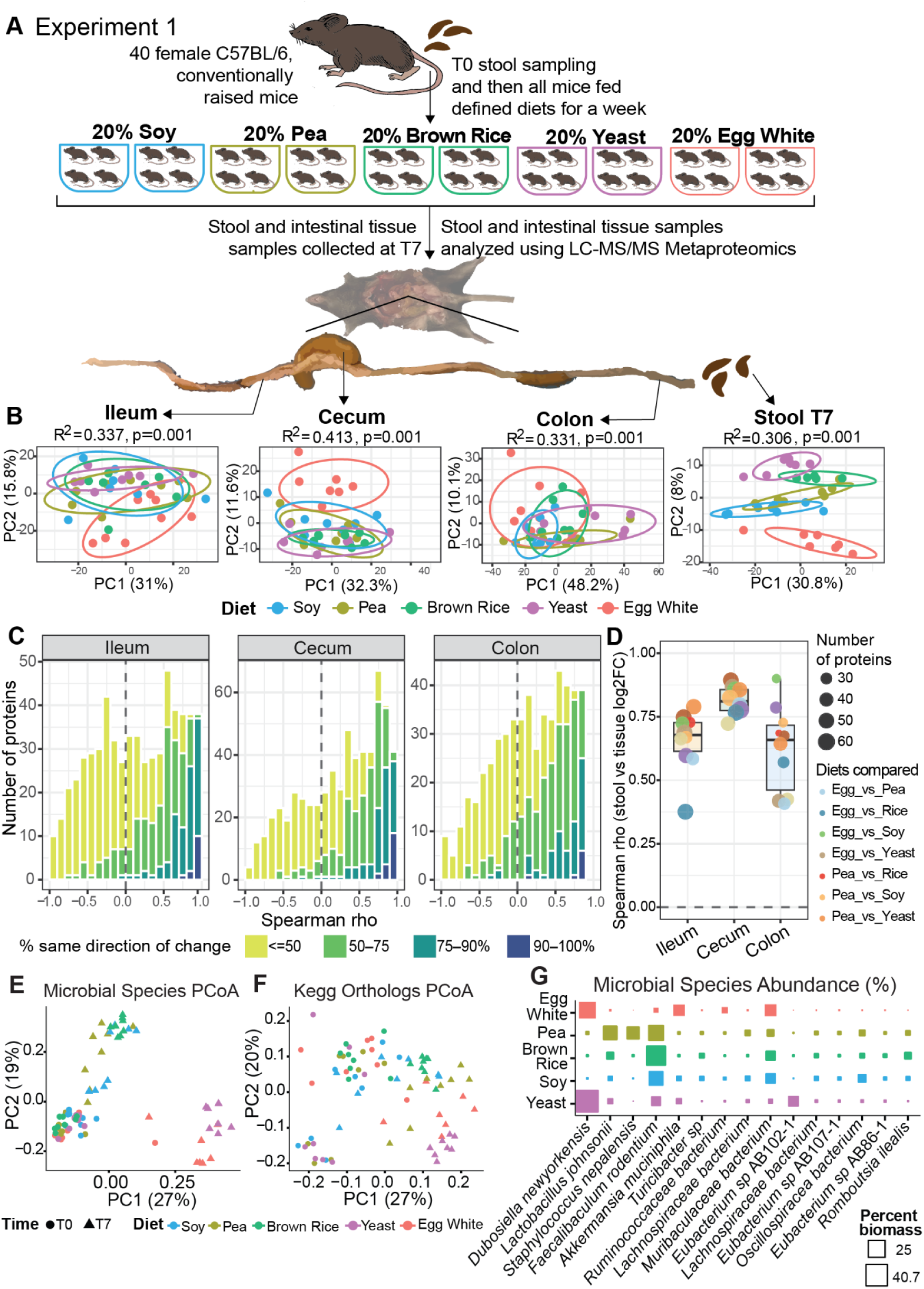
Stool and tissue analyses reveal that dietary protein source induces region-specific host responses in the intestinal tract, and most of these region-specific responses are captured in the stool. A) Experimental design schematic for Experiment 1. Mice were fed defined diets containing purified dietary protein from soy, pea, brown rice, yeast, or egg white for a week (n=8 in 2 cages per diet). At the end of the week, at T7, stool and tissue samples were collected and analyzed using LC-MS/MS metaproteomics (all conditions had n=8 replicates, except egg white, which had n=7 for the day 7 stool samples). B) PCA plots showing differences in the host proteomes of mice fed different dietary protein sources in the ileum, cecum, colon, and stool. Diet effects on the host proteome were determined using PERMANOVA as implemented in the vegan R package based on Euclidean distances with diet as the main predictor variable over 999 permutations, and the resulting R2 and p values are reported for each tissue^35,36^. Dispersion homogeneity was confirmed in each tissue using betadisper (not significant in any tissue or stool) to ensure that PERMANOVA results reflected diet differences and not within-group dispersion^37^. Ellipses are based on an 80% multivariate t-distribution. C) Histogram showing the distribution of per-protein concordance in host protein responses between each of the tissues (ileum, cecum and colon) and the stool. For each protein, Spearman’s rank correlation rho was determined between its stool and tissue log2 fold changes across all pairwise dietary comparisons (10 comparisons) to quantify how well region-specific host intestinal responses are captured in the stool. Bar colors represent the percentage level of agreement in the direction of protein response. Bars to the right of the dashed line in light and dark green represent proteins with high rho correlation values and high directional agreement (>=75%) between tissue and stool responses. Only proteins present in at least 6 pairwise dietary comparisons (643 distinct proteins across all tissues) were included in the correlation analysis, as estimating correlation based on fewer than 6 data points can give misleading results. D) Boxplot showing Spearman’s rank correlation between stool and tissue log2 fold changes across all proteins for each pairwise-dietary comparison in the ileum, cecum and colon. Each point represents a diet comparison. Only proteins with a |log2 fold change| >= 1 in both the stool and tissue were used for the correlation analysis used in the boxplot to determine concordance in proteins that are responding to protein source. E) and F) Bray-Curtis dissimilarity-based PCoA plots showing dietary protein source-driven differences in E) gut microbiota composition based on summed abundance of proteins assigned to MAG-based species groups^13^ and F) protein abundances summed into KEGG Orthologous groups to observe shifts in microbiota function (Diet R²=0.172, compared to T0 baseline R²=0.205, Diet:Baseline R²=0.142; all p=0.001). G) Square matrix plot showing the mean relative abundance of the most abundant microbial species (MAGs) across all dietary protein sources, where each species has an average relative abundance of at least 5% in at least one of the protein sources^38^. Data used in this figure are available in Supplementary Table 1.

To determine how well these site-specific diet-induced changes in the abundance of each protein were reflected in the stool proteome irrespective of dietary comparisons, we compared the direction and magnitude of changes in host protein abundances in the ileum, cecum, and colon to those in stool across all pairwise dietary comparisons (10 comparisons). For each protein changing in at least 6 pairwise dietary comparisons in the stool and tissue, which is equivalent to detection in at least 4 out of 5 diet groups, we quantified the percentage of pairwise dietary comparisons in which the fold change in the tissue and stool had the same direction (Fig. 1C). Of the 643 distinct proteins analyzed between all three gut sites and stool, an average of 50% showed directional concordance with stool in more than half of all the pairwise dietary comparisons. An average of 15% had strong directional agreement in at least 75% of dietary comparisons and 3% had strong directional agreement in at least 90% of dietary comparisons.

To further assess the concordance in the magnitude of change of each protein between the tissue and the stool, we quantified the rank correlation (Spearman’s rho) between diet-driven log2 fold changes in protein abundance in the tissue and stool across all pairwise dietary comparisons (Fig. 1C). Across all three gut sites, an average of 36% of proteins had at least a moderate positive correlation (rho >0.5), and an average of 21% had a strong positive correlation (rho >0.7) between tissue and stool responses. Host responses captured in stool had the highest concordance with the cecum, with 26.5% of proteins strongly correlating (rho >0.7) with host responses captured in the stool, followed by the colon (19%) and the ileum (18.8%).

We also assessed concordance between global tissue and stool responses for each pairwise dietary comparison to determine whether global diet-induced tissue responses at specific sites in the gut are captured in the stool. In each pairwise dietary comparison, we quantified the rank correlation between the log2 fold change of all proteins in both the stool and the tissue. Across all pairwise diet comparisons in each tissue, the magnitude of the host response in the stool was strongly positively correlated with that of the underlying tissue, with the cecum having the highest correlation across all dietary comparisons, followed by the ileum and the colon, which showed more variable responses (Fig. 1D). Overall, these results show the significant impact of dietary protein source on the host across the gut and that stool is able to capture many of these host responses occurring in specific sites along the intestinal tract.

### Dietary protein source is associated with shifts in the gut microbiota

We also evaluated shifts in gut microbial composition and function associated with dietary protein sources. Microbial community composition was significantly different between dietary protein sources and compared to the T0 baseline (Diet R²=0.263, compared to T0 baseline R²=0.222, Diet:Baseline R²=0.168; all p=0.001) (Fig. 1E). Across all treatment groups, microbial communities clustered together at the baseline (T0), where mice were fed the same standard chow. They diverged significantly from the baseline at day 7 (T7) and separated by dietary protein source. Egg white and yeast groups diverged the most from the baseline and from the other groups, while the Soy group diverged the least from the baseline. At the taxonomic level, these changes were driven by changes in abundance of specific taxa. In the egg white protein and yeast protein groups, *Dubosiella newyorkensis* was enriched, while *Faecalibacterium rodentium* was depleted compared to the other diet conditions (Fig. 1F). Additionally, *Akkermansia muciniphila* was more abundant in the egg white protein group compared to all other groups, while *Lactobacillus johnsonii* and *Staphylococcus nepalensis* were more abundant in the pea protein group compared to all other groups.

Dietary protein source further impacted gut microbial function by driving differences between the KEGG Ortholog (KO) profiles of dietary groups. As with gut microbial composition, both dietary protein source and comparison to the baseline significantly impacted microbial functional profiles (Diet R²=0.172, compared to T0 baseline R²=0.205, Diet:Baseline R²=0.142; all p=0.001) (Fig. 1F). Functional profiles for the yeast, egg whites, and brown rice protein groups diverged the most from the baseline and from other groups. Overall, these results show the significant impact of dietary protein source on gut microbial composition and function.

### Dietary protein source impacts host physiology through microbiota-dependent and independent effects

To determine which of the impacts of dietary protein source on gut physiology are directly driven by protein source and which are mediated by the gut microbiota’s response to dietary protein source, we compared the composition of host proteins detected in stool samples of germ-free and conventional mice fed purified dietary protein from casein, egg whites, peas, brown rice, soy and yeast (Fig. 2, Supp tables 2 and 3). We extracted the stool host proteome from metaproteomic data previously used to describe microbiota responses to dietary protein source and the fate of dietary protein in the gut^13,14^ (Fig. 2A). We found that the presence of a gut microbiota significantly impacted the composition of the stool host proteome across all eight diets (PERMANOVA: R^2^=0.382, p<0.05) (Fig. 2B). PCA analysis showed that the stool host proteomes of conventional and germ-free mice separated along the first component (Fig. 2B). Within the germ-free and conventional groups, the host proteome was significantly different across all six dietary protein sources (Fig. 2C and D). The male and female groups clustered separately only in the conventional mice fed the yeast, egg white, and brown rice dietary proteins and this difference was associated with differences in gut microbiota composition characterized in a previous study^13^ (Suppl. Fig. 2B). No separation between the male and female groups was observed in any of the germ-free samples (Suppl. Fig. 2A). Additionally the 20% and 40% soy and casein groups did not separate hence for all analyses downstream we focused only on the host responses in the 20% diets.

**Figure 2.**
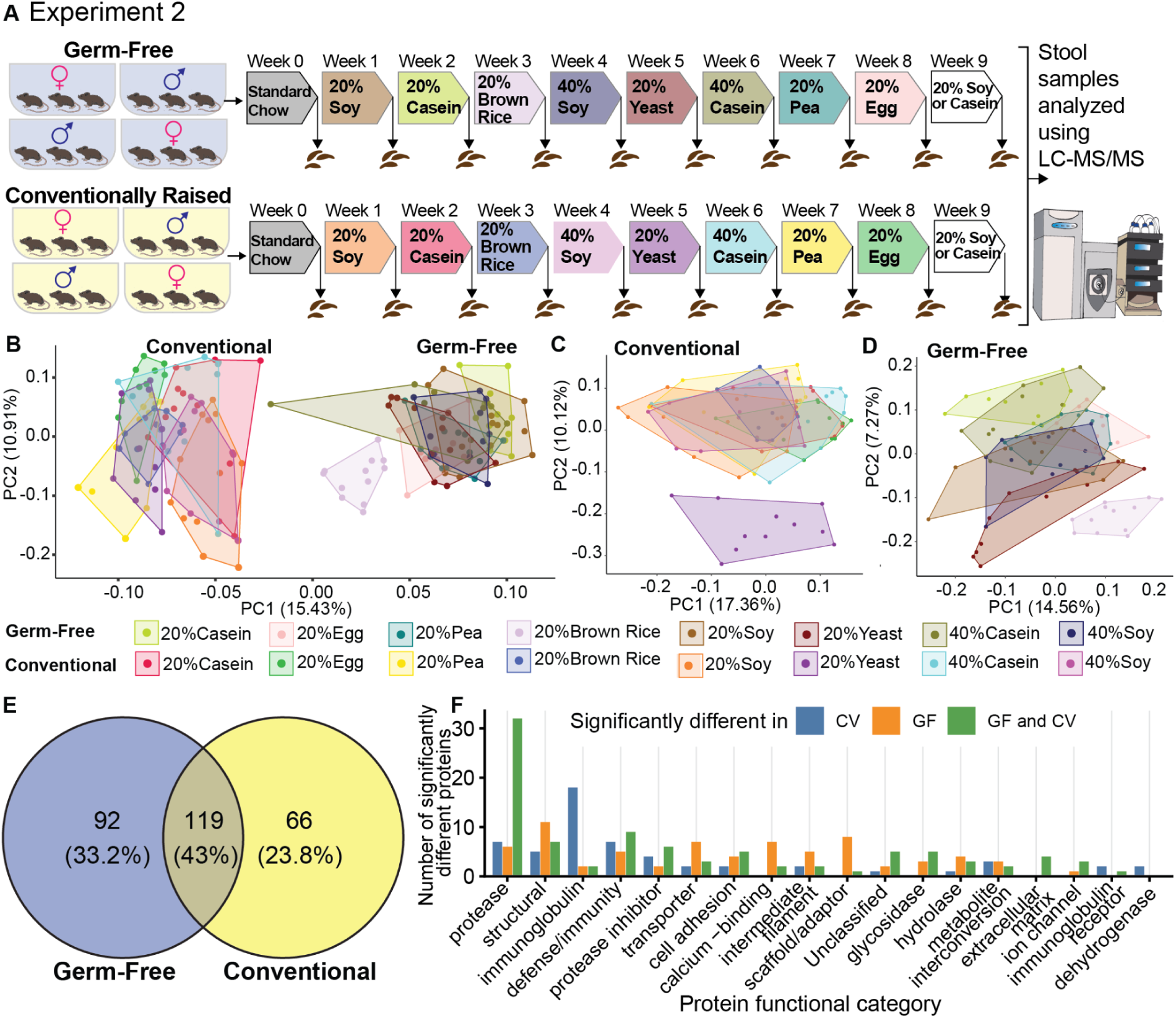
Dietary protein source impacts stool host proteome in both the presence and absence of the gut microbiota. A) Experimental design schematic for Experiment 2. Germ-free and conventionally raised mice were fed defined diets containing purified dietary protein from casein, soy, peas, brown rice, egg whites, and yeast for one week, and stool samples were collected for LC-MS/MS based metaproteomics. B-D) PCA plots showing compositional differences between the stool host proteomes of germ-free and conventional mice fed defined diets with a single protein source, (20% casein (n=11), 20% egg white (n=10), 20% pea (n=10), 20% brown rice (n=11), 20% soy (n=12), 20% yeast (n=12), 40% casein (n=10) or 40% soy (n=10)). E) Venn diagram showing the number and percentage of host proteins significantly different in at least one pairwise comparison across the eight diets in the Germ-Free group only, the Conventional group only, and in both the Germ-Free and Conventional groups. F) Barplot showing the number of significantly different proteins in various functional categories in the Germ-Free group only (orange), the Conventional group only (blue), and in both the Germ-Free and Conventional groups (green). Only the top 10 categories containing the most differentially abundant proteins across the 3 groups are shown. The data underlying this figure are available in Supplementary Table 2 and Supplementary Table 3.

We found that 277 host proteins were significantly different in at least one pairwise comparison of different dietary protein sources, with pairwise comparisons done separately for germ-free and conventional mice. We used the comparison between germ-free and conventional mice to classify these host responses to dietary protein source as microbiota-dependent or microbiota-independent. Host responses that overlapped between germ-free and conventional mice were classified as microbiota-independent since they occurred regardless of the presence of the microbiota and responses found only in conventional mice were classified as microbiota dependent (Fig. 2E). Host responses unique to germ-free mice were not included in the downstream analysis as they are difficult to interpret in the context of the well characterized developmental and physiological differences observed in germ-free mice^39,40^. Of the 185 proteins that responded to dietary protein source in conventional mice, a large proportion, 64% (119 out of 185), also responded in germ-free mice, indicating microbiota-independent host responses to dietary protein source (Fig. 2E). The remaining 36% (66 out of 185) of proteins responded to dietary protein source only in conventional mice, indicating microbiota-dependent responses.

We further determined the main functional categories of the differentially abundant host proteins using Gene Ontology and PANTHER protein classification. We found that the majority of proteins that were significant in both germ-free and conventional mice were classified as proteases, defense/immunity-related proteins, structural proteins, protease inhibitors, and cell adhesion molecules (Fig. 2F). Host proteins differentially abundant in only the germ-free group mostly fell in the structural, scaffold/adaptor, transporter, calcium-binding, and protease categories. Host proteins differentially abundant across different diets in only the conventional group fell in the immunoglobulin, protease, defense/immunity, structural, and protease inhibitor categories. Overall, we observed differences in protein categories in germ-free and conventional mice across the dietary protein sources. To explore this further, we focused on specific proteins within the categories that had the greatest number of differentially abundant proteins. These included proteases, protease inhibitors, immune proteins, and proteins involved in intestinal epithelial maintenance or barrier function, which were distributed across the structural, cell adhesion, and defense categories. Overall, these results show that dietary protein source impacts several aspects of gut physiology both in the presence and absence of the gut microbiota. More in-depth comparisons of proteins across dietary protein sources in these categories are presented below.

### Host proteases and protease inhibitors differentially respond to changes in dietary protein source in both the presence and absence of gut microbes

Since host proteases and protease inhibitors were particularly numerous among proteins differentially abundant between dietary protein sources in both the germ-free and conventional groups (45 in total), we analyzed these proteins in detail. We focused on proteins that responded to multiple protein sources by identifying host proteases and protease inhibitors that were significantly enriched or depleted in a dietary protein source compared to at least four other dietary protein sources.

We identified 10 different digestive proteases that were significantly higher in abundance in the egg white or brown rice protein groups in just the conventional mice or in both germ-free and conventional mice (Fig. 3). Proteases that responded to dietary protein source in both germ-free and conventional conditions were considered microbiota-independent responses while the ones that responded in only the conventional condition were considered to be microbiota-dependent responses. Both serine protease 3 (PRSS3) and trypsin 10 (TRY10) were significantly higher in the egg white protein group compared to all other diets but only in conventional mice, indicating that this was a microbiota-dependent response. However, trypsin 3B (PRSS3B) was higher in the egg white protein group in both conventional and germ-free mice, indicating that this was a microbiota-independent response. Similarly, carboxypeptidase A1 (CPA1), A2 (CPA2), and B (CPB1), and chymotrypsin-like protease (CTRL) were significantly higher in the brown rice protein group compared to other groups in both conventional and germ-free mice, indicating that this was a microbiota-independent response. Chymotrypsin-like elastase family members (CELA1 and CELA2A) were significantly higher in the brown rice protein group in conventional mice, but only CELA1 was significantly higher in the brown rice protein group in germ-free mice, indicating a microbiota-independent response for CELA1 only. Chymotrypsinogen B (CTRB1) was higher in both egg white and brown rice protein groups in both conventional and germ-free mice, indicating a microbiota-independent response. These proteases are released into the small intestine as part of pancreatic secretions for digestion^41^.

**Figure 3.**
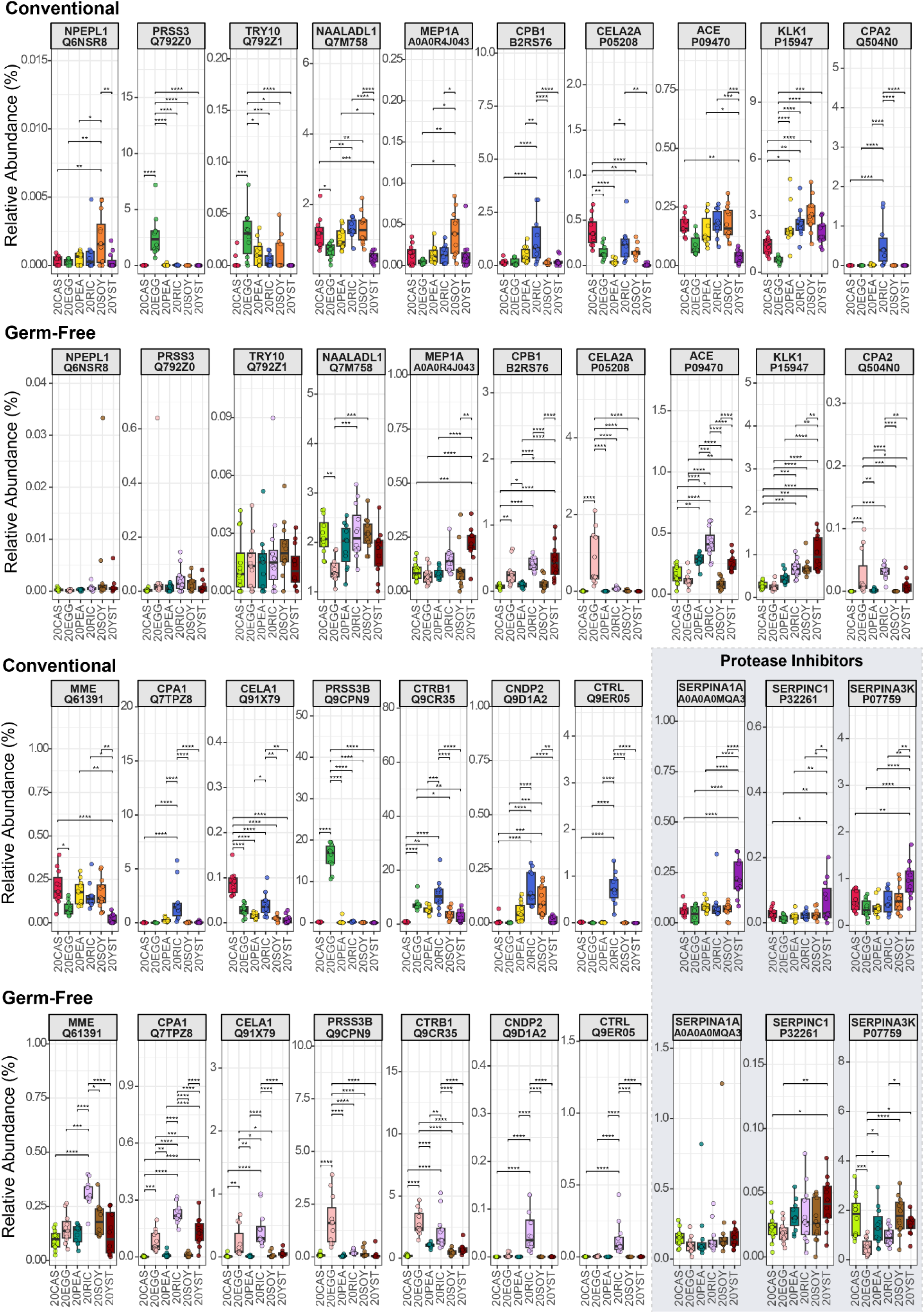
Dietary protein source impacts the abundance of host proteases and protease inhibitors in both conventional and germ-free mice. Boxplots showing the relative abundance (orgTSS%) of differentially abundant host proteases and protease inhibitors in stool of Conventional and Germ-free mice fed defined diets containing 20% protein (Fig. 2A) from casein (20CAS), egg white (20EGG), pea (20PEA), brown rice (20RIC), soy (20SOY) and yeast (20YST). The header strip includes the protein name and its UniProt accession. The shaded box differentiates the protease inhibitors from the proteases. Statistical significance, determined by ANOVA followed by Tukey’s HSD post hoc test, is indicated by horizontal lines and asterisks (✱= p<0.05, ✱✱ = p<0.01, ✱✱✱ = p<0.001, ✱✱✱✱ = p<0.0001). Proteases and protease inhibitors that were considered diet responders and significantly different in at least 4 pairwise comparisons per reference diet (e.g., at least 4 significant comparisons with egg white as reference, at least 4 with brown rice as reference, and so on) in conventional mice (conventional only and overlap with germ-free region of Venn diagram in Fig 2E) are shown. The complete underlying data set can be found in Supplemental Table 2.

Beyond digestive proteases, we identified tissue proteases and metalloproteases that responded to changes in dietary protein source. Probable aminopeptidase NPEPL1 was significantly higher in conventional mice fed soy protein only, indicating a microbiota-dependent response. For other proteases, the presence of the microbiota altered which dietary protein source the proteases responded to. These included aminopeptidase NAALADL1, which was lower in both egg white and yeast protein groups in conventional mice, but only lower in the egg white protein diet in the germ-free condition. Metalloprotease meprin 1A (MEP1A) was significantly higher in the soy protein group in conventional mice, but higher in the yeast protein group in germ-free mice. Peptidases angiotensin-converting enzyme (ACE) and neprilysin (MME) were significantly lower in conventional mice fed yeast protein compared to other diets, while they were highest in the brown rice protein group in germ-free mice. Kallikrein-1 (KLK1), a serine protease part of the kallikrein-kinin system, was lower in the egg white protein diet compared to other diets in both conventional and germ-free mice, indicating a microbiota-independent response. Cytosolic non-specific dipeptidase (CNDP2) was higher in the brown rice protein group in both conventional and germ-free mice, indicating a microbiota-independent response as well.

In addition to proteases, we also found that the abundance of protease inhibitors changed in response to dietary protein source (Fig. 3). Alpha-1-antitrypsin 1 (SERPINA1A), antithrombin-III (SERPINC1), and serine protease inhibitor A3K (SPA3K) were all significantly higher in the yeast protein group in conventional mice. However, only SERPINA1A responded in a microbiota-dependent manner. SPA3K shifted its response in the absence of the gut microbiota and was lowest in germ-free mice fed egg white protein. Overall, our results show that most proteases respond to dietary protein source independent of the gut microbiota in both germ-free and conventional conditions and that dietary protein sources shown to have lower digestive efficiency have higher abundance of digestive proteases.^14^

### Yeast protein induces an antimicrobial immune response in the gut, with REG3 lectins responding independently of the gut microbiota

Among the stool host proteins that significantly changed due to dietary protein source in Experiment 2 (Fig. 2F), we identified eight host proteins associated with the host’s antimicrobial immune response that were significantly enriched or depleted in one protein source relative to four other protein sources in conventional mice or in both germ-free and conventional mice. Several of these immune proteins responded only in the conventional condition, indicating a dependence on microbial interactions with the host and the diet. These microbiota-dependent diet-responder proteins were particularly elevated following the yeast protein diet. Immunoglobulin alpha chain C (IGHA), prolactin-inducible protein homolog (PIP), and calprotectin subunit A9 (S100A9) were all significantly enriched in conventional mice fed the yeast protein diet, but did not change in abundance in the germ-free group (Fig. 4). These proteins play immunomodulatory and antimicrobial roles in the gut and calprotectin subunit A9 (S100A9), is a clinical marker of inflammation^42,43^. However, Zymogen granule protein 16 (ZG16), an antimicrobial lectin-like protein produced by goblet cells that helps maintain the intestinal barrier by limiting bacterial translocation^44^, was significantly lower in conventional mice fed egg white protein, but did not change in the germ-free mice.

**Figure 4.**
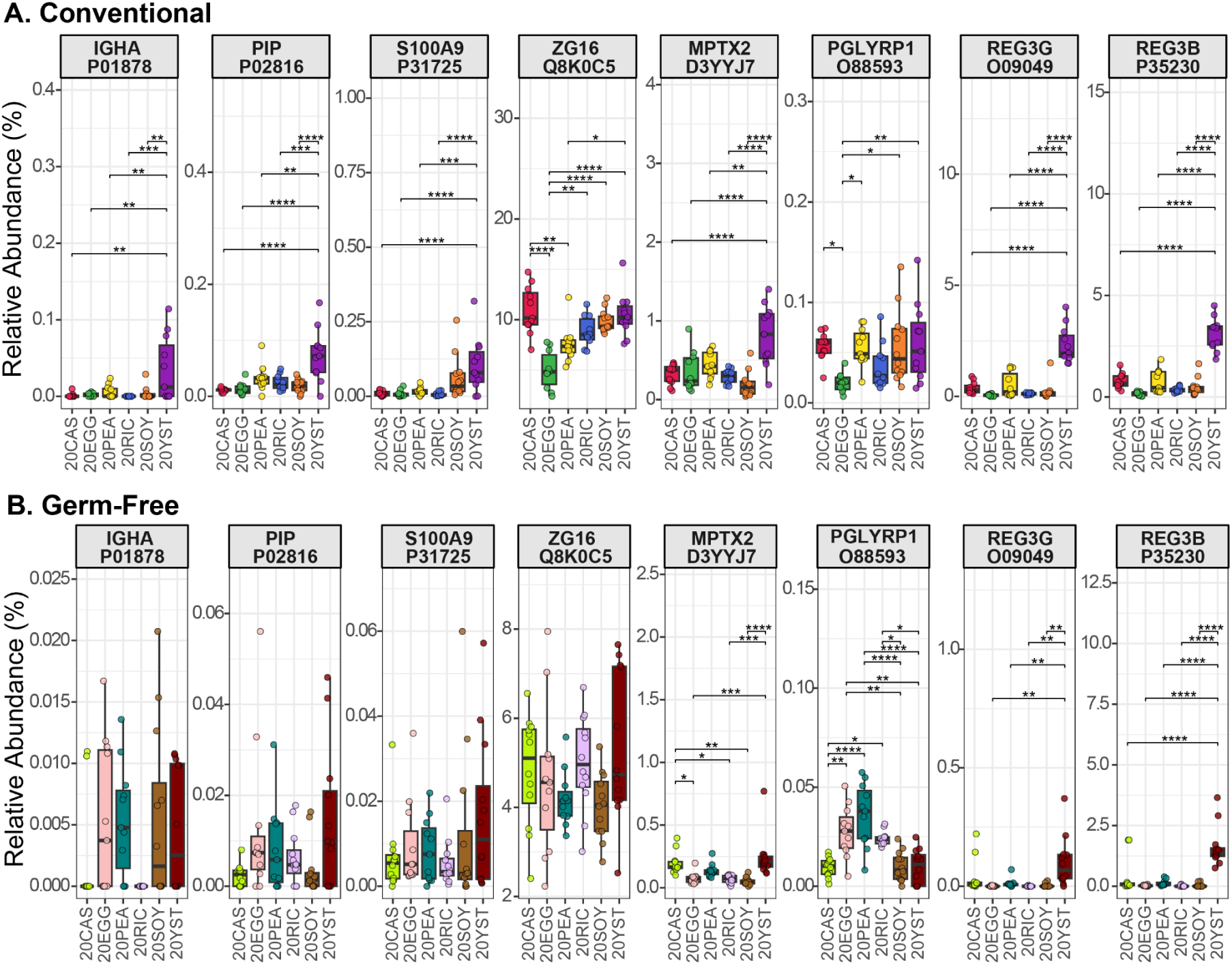
Immune proteins with antimicrobial activity change in abundance in response to different dietary protein sources regardless of microbial status. Boxplots showing the relative abundance (orgTSS%) of differentially abundant immune proteins in stool of A) Conventional and B) Germ-free mice fed defined diets containing 20% protein (Fig. 2A) from casein (20CAS), egg white (20EGG), pea (20PEA), brown rice (20RIC), soy (20SOY) and yeast (20YST). The header strip includes the protein name and its UniProt accession. Statistical significance as determined by ANOVA followed by Tukey’s HSD post hoc test is indicated by horizontal lines and asterisks (✱= p<0.05, ✱✱ = p<0.01, ✱✱✱ = p<0.001, ✱✱✱✱ = p<0.0001). Immune proteins that were considered diet responders and significantly different in at least 4 pairwise comparisons per reference diet in conventional mice (conventional only and overlap with germ-free region of Venn diagram in Fig 2E) are shown. The complete underlying data set can be found in Supplemental Table 2.

Alternatively, other immune proteins changed both in the presence and absence of the gut microbiota, demonstrating a microbiota-independent response. These included pentraxin family member protein (MPTX2), a Paneth cell marker that plays an important role in regulating the composition of the gut microbiota, which was significantly higher in both conventional and germ-free mice fed yeast protein compared to other protein sources^45,46^. Peptidoglycan recognition protein 1 (PGLYRP1), an antimicrobial pattern recognition receptor, was significantly lower in conventional mice fed egg white protein, but in germ-free mice it was significantly higher in the egg white, pea and brown rice protein groups. Antimicrobial proteins Reg3γ (REG3G) and β (REG3B) were significantly higher in mice fed the yeast protein compared to all other dietary protein sources in both the germ-free and conventional groups^47^. These results show that yeast protein is associated with an increase in the abundance of several antimicrobial and immune proteins only in the presence of the gut microbiota. However, REG3 lectins specifically are impacted independent of the presence of a microbiota.

### Intestinal barrier is impacted by dietary protein source both directly and via diet-microbiota interactions

We identified 11 proteins associated with the maintenance of the intestinal epithelium that were significantly enriched or depleted in one protein source relative to four other protein sources in conventional mice or in both germ-free and conventional mice. Protein S100A6 (calcyclin), which is associated with epithelial proliferation in the gut, was significantly higher in only conventional mice fed soy protein, indicating a microbiota-dependent response (Fig. 5). The rest of the proteins responded in both conventional and germ-free mice to dietary protein source, and some had consistent microbiota-independent responses to specific protein sources. Carcinoembryogenic antigen-related cell adhesion molecule 1 (CEACAM1), which plays a role in cell adhesion in the intestinal epithelium and modulates mucosal immune response, was significantly higher in conventional mice fed soy and yeast protein. In germ-free mice, CEACAM1 was significantly higher in both the yeast protein and casein protein groups compared to the egg white and brown rice protein groups. Cadherin-related family protein 5 (CDHR5), which supports gut barrier integrity by forming intermicrovillar adhesion complexes (IMACs), was significantly lower in the egg white and yeast protein groups and higher in the casein protein group in both conventional and germ-free mice, indicating a microbiota-independent response^48^.

**Figure 5.**
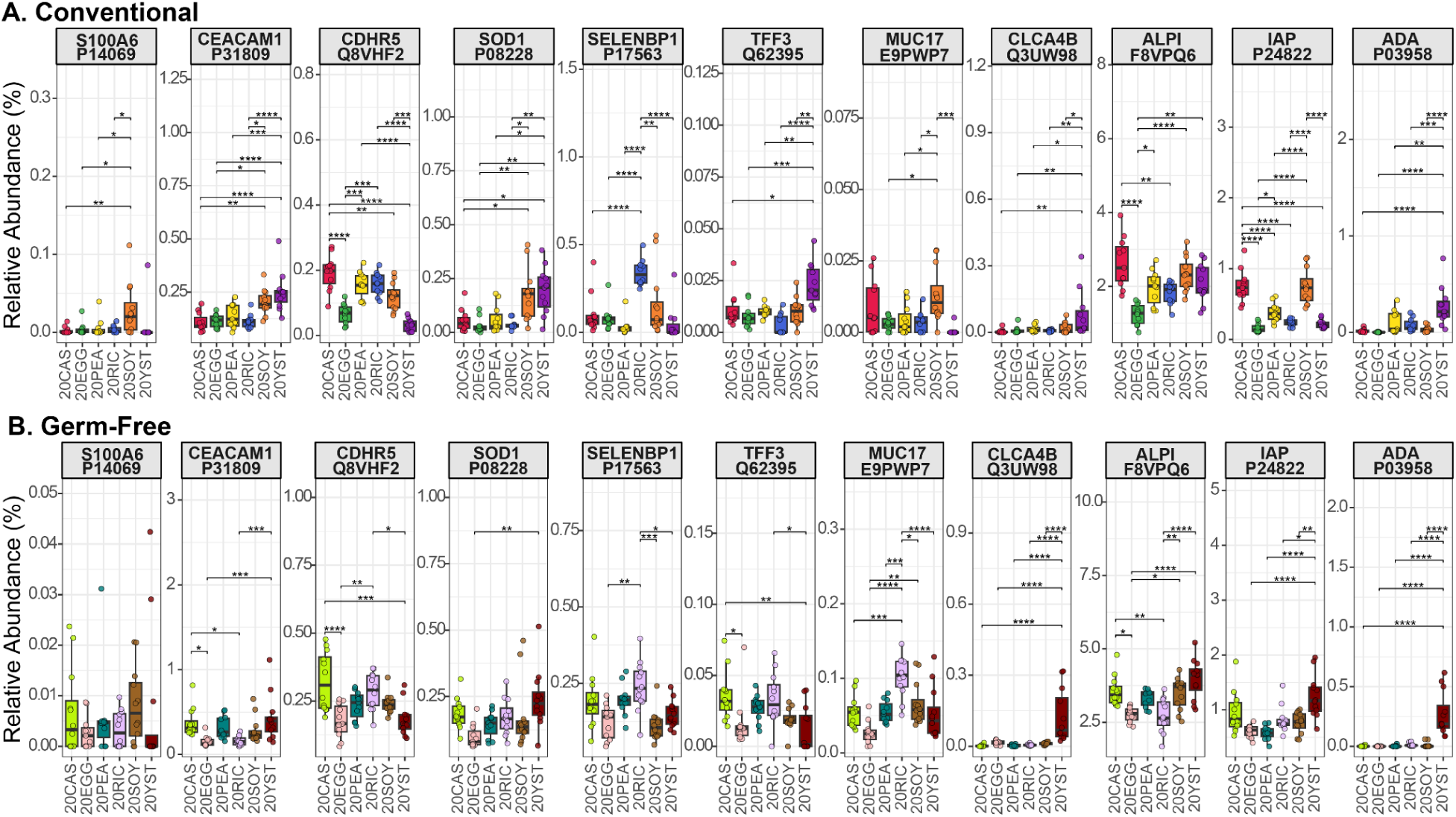
Proteins related to barrier function and maintenance of the intestinal epithelium change in abundance in response to different dietary protein sources. Boxplots showing the relative abundance (orgTSS%) of differentially abundant barrier function and intestinal epithelial maintenance proteins in stool of A) Conventional and B) Germ-free mice fed defined diets containing 20% protein (Fig. 2A) from casein (20CAS), egg white (20EGG), pea (20PEA), brown rice (20RIC), soy (20SOY) and yeast (20YST). The header strip includes the protein name and its UniProt accession. Statistical significance, determined by ANOVA followed by Tukey’s HSD post hoc test, is indicated by horizontal lines and asterisks (✱= p<0.05, ✱✱ = p<0.01, ✱✱✱ = p<0.001, ✱✱✱✱ = p<0.0001). Proteins that were considered diet responders and significantly different in at least 4 pairwise comparisons per reference diet (e.g., at least 4 significant comparisons with egg white as reference, at least 4 with brown rice as reference, and so on) in conventional mice (conventional only and overlap with germ-free region of Venn diagram in Fig 2E) are shown. The complete underlying data set can be found in Supplemental Table 2.

Superoxide dismutase (SOD1), an antioxidant that protects the gut epithelium against oxidative stress, was significantly higher in conventional mice fed soy and yeast protein compared to most other groups. In germ-free mice, SOD1 was significantly higher in the yeast protein group compared to egg white protein only. Trefoil factor 3 (TFF3), a goblet cell marker protein, was higher in conventional mice fed yeast protein, while in the germ-free group, it was lower in the yeast protein group compared to the casein and brown rice protein groups^49^. Mucin 17 (MUC17), a transmembrane protein contributing to the mucus layer produced by mature epithelial cells, was significantly higher in conventional mice fed soy protein, but in germ-free mice it was higher in the brown rice group compared to all other groups^50^. Calcium-activated chloride channel regulator 4B (CLCA4B), a regulator of gut mucus homeostasis that is upregulated in the colon in gut inflammation^51^, was significantly higher in both conventional and germ-free mice fed yeast protein, indicating a microbiota-independent response. We detected two forms of intestinal-type alkaline phosphatase, which dephosphorylates lipopolysaccharide and plays a role in maintaining gut barrier function ^52,53^: ALPI, which is expressed throughout the gut, and IAP, which is expressed mainly in the small intestine^54^. ALPI was significantly lower in the egg white group in both conventional and germ-free mice. IAP was significantly higher in conventional mice fed casein and soy protein, but in germ-free mice it was higher in the yeast protein group. Adenosine deaminase (ADA), which is involved with the development of Peyer’s patches and is elevated in gut inflammation, was significantly higher in both conventional and germ-free mice fed yeast protein.

To further investigate the impact of dietary protein source on the mucus barrier, we quantified shifts in the abundance of host proteins associated with the mucus layer in stool samples from the conventional mice in Experiment 1. We also measured the thickness of the mucus layer in the colon tissue of these mice using Alcian Blue staining (Fig. 6). We found that conventional mice fed egg white protein showed significantly reduced summed relative abundance of mucus proteins compared to the other groups (Fig. 6A). These host responses in stool paralleled tissue level responses where the egg white protein group had significantly reduced mucus layer thickness compared to the brown rice and soy protein groups (Fig. 6B and C). To determine whether these changes in the host were associated with changes in specific microbial functions in Experiment 2, we quantified the impact of dietary protein source on the abundance of microbial proteins that are associated with mucin degradation in the gut, including glycoside hydrolases (GH) that degrade mucin glycosylations and enzymes with the capacity to release and consume the components of mucin glycosylations (sialic acid, acetylglucosamine, and fucose). The egg white protein group had a significantly higher abundance of these mucin degradation-associated proteins (Fig. 6D).

**Figure 6.**
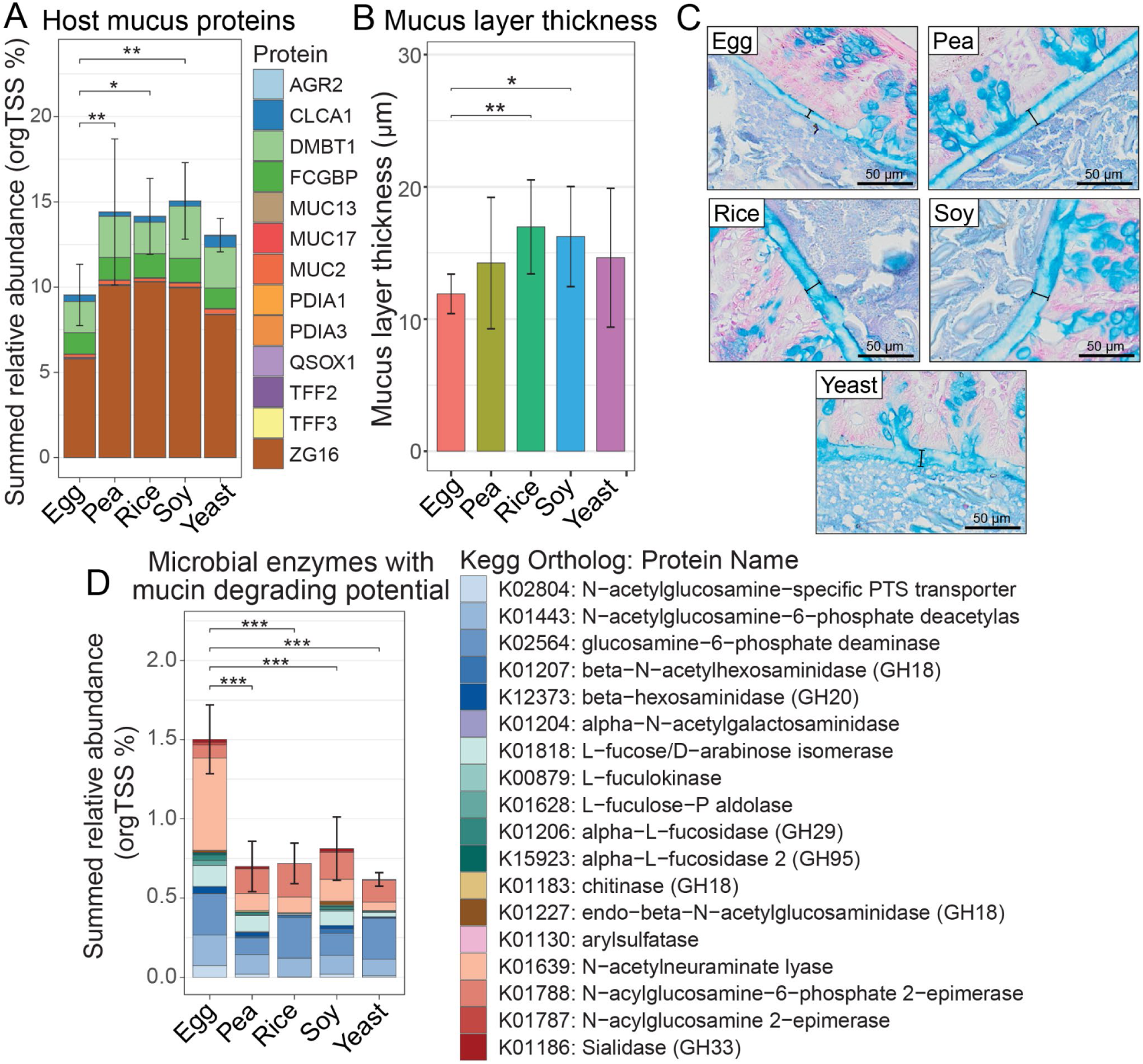
Dietary protein source impacts mucus layer thickness, as well as the abundance of host mucus-associated proteins and microbial mucus-degrading enzymes (Experiment 1, Fig. 1A). A) Summed relative abundance of host proteins associated with the mucus layer. Each stack is the mean relative abundance of a protein, and the stacked bars represent the summed mean relative abundance of all mucus-related proteins in each diet. B) Colonic mucus layer thickness was measured in mice fed diets containing protein isolates from egg whites, pea, soy, yeast, and brown rice. C) Representative images showing alcian blue staining of the mucus layer in colons of mice fed different dietary protein sources. Brackets in images indicate the mucus layer. D) Summed relative abundance of microbial enzymes associated with mucus degradation in the stool. Statistical significance, determined by a pairwise t-test followed by Benjamini-Hochberg correction for multiple tests, is indicated by horizontal lines and asterisks (✱= p<0.05, ✱✱ = p<0.01, ✱✱✱ = p<0.001, ✱✱✱✱ = p<0.0001). Bars represent standard deviation from the group means. All data in this figure are from conventionally raised mice in Experiment 1 (Fig. 1A).

These results are consistent with our previous microbiota-focused analysis of Experiment 2, in which egg white protein similarly promoted the abundance of these mucin degradation-associated microbial proteins, a finding we further validated *in vitro*^13^. Interestingly, the summed abundance of stool host mucus proteins was also significantly lower in the egg white protein group than the brown rice protein group in germ-free mice from Experiment 2, indicating that egg white protein can impact the mucus barrier even in the absence of the gut microbiota (Supp. Fig. 2). However, this difference was smaller in magnitude in the germ-free mice compared to the corresponding change in the conventional mice. Overall, these results show that dietary protein source impacts the gut barrier, altering the thickness of the mucus layer in tissue and the abundance of gut microbial enzymes associated with degrading the mucus layer.

### Dietary protein source effects on the host and microbiome translate to humans

Since the effects of dietary protein source we observed in mice were based on defined diets consisting of purified dietary components, we sought to determine the extent to which these dietary protein source effects on the host and gut microbiota translate to humans eating a complex diet. For this, we compared how the host and microbial components of the stool metaproteome shift in response to egg white and pea protein consumption in a previously described randomized crossover controlled feeding trial in healthy humans^55^. Participants in this study ate a diet where 70% of their protein was from either an egg white or pea protein supplement (Fig. 7A). We re-analyzed these samples using data-independent acquisition (DIA) mass spectrometry and a metagenome-assembled genomes (MAGs) based sample-matched metagenomic database.

**Figure 7.**
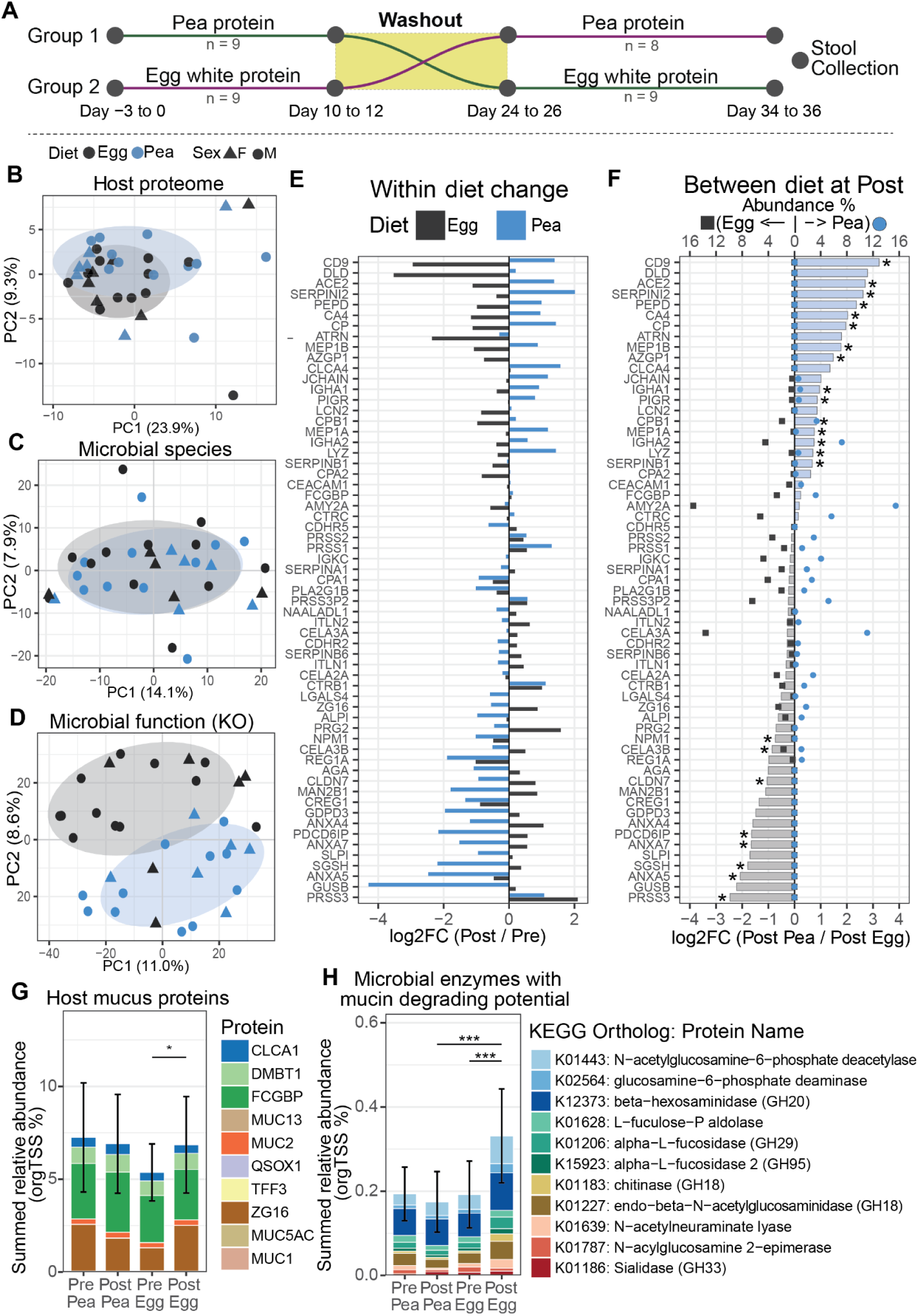
Dietary protein source reshapes host responses and microbial mucin-degrading potential in human stool metaproteome. A) Schematic for the crossover randomized controlled feeding trial. B-D) PCA plots showing post dietary intervention differences in B) the stool host proteome, C) microbiota composition based on summed abundance of proteins assigned to MAG-based species groups, and D) microbiota function based on protein abundances summed into KEGG Orthologs. Samples from subjects fed egg white protein are colored black and samples from subjects fed pea protein are colored blue. Ellipses are based on a 68% multivariate t-distribution. E) Barplot depicting within-diet log2 fold change (log2FC) in the abundance of diet responder proteins between pre and post diet intervention (i.e., comparing the baseline values pre-diet to the post intervention values for the same individuals in a paired manner). Post vs pre log2FC bars are black for egg white and blue for pea. F) Barplot overlaid with abundance data showing the between-diet paired log2FC in the abundance of the same diet responder proteins as shown in B between post pea and post egg white protein diets. Bars are shaded blue where the protein was higher in the pea protein group and grey where protein was higher in the egg white protein group. Two scatter plots with a secondary x-axis on the top of the plot depict the mean post intervention relative abundance (orgTSS%) of proteins in the pea (blue circles, right of the 0) or egg white (black squares, left of the 0) protein groups. Statistical differences in F were determined using a paired t-test followed by a Benjamini-Hochberg correction for multiple comparisons, and asterisks denote proteins that were significantly different between the two diets at the post-intervention timepoint. Diet responder proteins shown in E and F are ordered by the magnitude of their log2FC between diets at post intervention and were selected based on the following criteria: 1) a significant diet effect in a linear model comparing pre vs post log2FC change between diets (difference between diets in the change induced from pre to post intervention); 2) a significant diet effect at post intervention between diets in an ANCOVA of post protein abundance adjusted for pre abundance with individual as a random intercept (difference between diets at post-intervention); or 3) a mean relative abundance (orgTSS%) of at least 1% in any diet or timepoint. G) Summed relative abundance of host proteins associated with the mucus layer detected in the human stool samples. Each stack is the mean relative abundance of a protein, and the stacked bars represent the summed mean relative abundance of all mucus-related proteins in each diet. F) Summed relative abundance of microbial enzymes associated with mucus degradation in human stool samples. Statistical significance in G and H was determined using a paired t-test followed by Benjamini-Hochberg correction for multiple tests, and is indicated by horizontal lines and asterisks (✱= p<0.05, ✱✱ = p<0.01, ✱✱✱ = p<0.001, ✱✱✱✱ = p<0.0001). Bars represent standard deviation from the group means.

In terms of the host response, our analysis revealed no significant differences in the stool host proteome of the two groups at baseline. However, both groups had a significant impact of protein source post dietary intervention (PERMANOVA on Euclidean distances: Diet R²=0.031, p=0.012; Sex R²=0.034, p=0.039). We further found that host proteins pertaining to different aspects of gut physiology changed in abundance in response to dietary protein source in human stool, as was previously observed in mice. These host proteins included digestive proteases such as trypsin and CELA3B, serine protease inhibitor SPI2, antimicrobial proteins such as IGHA and LYSC, and gut epithelial and barrier proteins such as tetraspanin and several annexins (Fig. 7E and F).

Several digestive proteases responded differentially to egg white and pea protein. Serine protease 3 (PRSS3) increased in response to dietary intervention in both pea and egg white protein groups, but the increase was larger in magnitude in response to egg white protein consumption. Chymotrypsin-like elastase family member 3B (CELA3B) increased from baseline in response to egg white protein and decreased in response to pea protein. CELA3B was also significantly higher in the egg white protein group post intervention compared to the pea protein group. Carboxypeptidase B (CPB1) was higher in the pea protein group compared to the egg white protein group post intervention. This difference in CPB1 between the two groups post intervention was largely due to an egg white protein driven decrease in its abundance from baseline rather than a large increase in response to pea protein.

Proteases not related to digestion and protease inhibitors also changed in response to dietary protein source. Meprin A (MEP1A) and B (MEP1B), subunits of the metallopeptidase meprin that plays a role in barrier maintenance, as well as Angiotensin-converting enzyme 2 (ACE2) and Xaa-Pro dipeptidase (PEPD), were all significantly higher in abundance in the pea protein group post intervention compared to the egg white protein group. The pea protein group had a significantly higher abundance of serine protease inhibitors such as serpin I2 (SERPINI2) and leukocyte elastase inhibitor (SERPINB1) compared to the egg white protein group post intervention. However, interestingly, the change in SERPINI2 abundance at post intervention was due to a pea protein driven increase from baseline, while the difference in SERPINB1 post intervention was due to an egg white protein driven decrease from baseline.

Host proteins related to the immune response and intestinal epithelial maintenance also changed in abundance in response to dietary protein source. Pea protein induced an increase in the abundance of antimicrobial immune response proteins immunoglobulin heavy constant alpha 1 (IGHA1) and 2 (IGHA2), polymeric immunoglobulin receptor (PIGR), which is involved in the secretion of IgA into the gut, and lysozyme C (LYZ). Tetraspanin (CD9) and carbonic anhydrase 4 (CA4) were also higher in the pea protein group post intervention compared to the egg white protein group. On the other hand, the egg white protein group had a higher abundance of annexin A5 (ANXA5) and A7 (ANXA7), claudin-7 (CLDN7), programmed cell death 6-interacting protein (PDCD6IP), N-sulphoglucosamine sulphohydrolase (SGSH) and nucleophosmin (NPM1) compared to the pea protein group post intervention.

To determine the effect of dietary protein source on the microbiota, the original analysis of this human crossover study used unassembled metagenomic reads and metaproteomes^55^ and found that while a few specific Firmicutes species were significantly more abundant in the pea protein group, neither alpha nor beta diversity differed significantly between the two diets. However, several microbial metabolic pathways differed significantly in abundance between the two dietary protein sources. In particular, amino acid biosynthesis was elevated in the pea protein group, while amino acid degradation enzymes were increased in the egg white protein group.

To assess if our re-analysis of the samples, using DIA analysis and a MAG-resolved approach, yielded similar results, we re-quantified microbiota composition by assessing per-species biomass by summing DIA-derived metaproteomic protein abundances by metagenome-assembled genomes (MAGs), which we assembled from metagenomic data^56^. We found that while there was no difference in taxonomic composition post-intervention (Fig. 7C), microbial functions in terms of protein abundances summed by KEGG Orthologs (KO) differed significantly (Fig. 7D).

In terms of microbial function, we specifically wanted to test whether the increase in microbial enzymes related to mucus degradation we previously observed in mice across two studies extended to humans. Our re-analysis revealed that the summed relative abundance of microbial enzymes related to the cleavage and consumption of the components of mucin glycans, e.g., fucose, acetylglucosamine, and sialic acid, was higher post egg white protein consumption compared to post pea protein consumption (Fig. 7H). These results show that egg white protein reproducibly promotes the abundance of enzymes that can degrade the glycan component of mucin across two distinct mouse experiments and one human experiment.

In terms of host mucus proteins, however, we found no difference between the pea and egg white protein groups post-intervention (Fig. 7G). Interestingly, we did find that host proteins associated with the mucus layer increased from pre- to post-intervention in the egg white group. This change was driven solely by an increase in the abundance of ZG16, which prevents bacterial penetration of the mucus layer^44^ (Supp. Fig. 3). Overall, our results show that the impacts of dietary protein source observed in mice extend, at least in part, to humans, in that protein source significantly alters the abundance of host proteolytic and antimicrobial proteins, and egg white protein is reproducibly associated with an increased abundance of microbial enzymes associated with mucin degradation.

## Discussion

Our goal in this study was to characterize the impact of dietary proteins from different sources on host responses in the gut. We aimed to decipher which effects were directly linked to diet-host interactions and which effects were mediated by the impact of dietary protein sources on the gut microbiota. Our results show that dietary protein source impacts gut physiology along the intestinal tract, with the majority of these responses captured in the stool. We further found that protein source effects on the host are only partially mediated by the gut microbiota as dietary protein source induced overlapping effects in both the presence and absence of the gut microbiota. These host responses were further associated with specific changes in gut microbial composition and function. Additionally, we found that the effects of dietary protein source on the host and gut microbiota extend to humans.

Our results show that dietary protein source reshaped the host proteome in the ileum, cecum, and colon, and these changes were largely reflected in the stool both in direction and magnitude (Fig. 1B-D). This concordance between site-specific host responses in the gut and host responses captured in the stool is a methodologically significant finding as it solidifies stool as a non-invasive readout of gut physiology, which is vital for measuring host responses in longitudinal studies. Additionally, both the concordant and discordant protein responses are informative as they carry different information specific to the location and response of the protein. For example, proteins secreted into the lumen, such as immunoglobulins and digestive enzymes, had high concordance between specific tissues and stool, while intracellular proteins had lower concordance or were discordant. Discordance of intracellular proteins may be due to changes that occur in cells after tissue is sloughed off from the epithelium. While it was beyond the scope of our study, further characterization of how individual protein or pathway responses in specific tissue sites correlate to stool responses could be a useful resource for mapping and measuring specific gut responses non-invasively in the stool, especially for clinical applications.

We evaluated which dietary protein source-induced host responses were mediated by the interactions of the diet with the gut microbiota and found that out of the 185 proteins that changed in response to dietary protein source in conventional mice, 119 (64%) also changed in the germ-free condition indicating that a majority of these protein source induced host responses are not dependent on the microbiota (Fig. 2F). We did not focus on germ-free only responses because germ-free mice have well characterized developmental and physiological abnormalities^39,40^ making it difficult to differentiate which of the germ-free specific responses were due to the diet versus which ones were due to germ-free physiology. The most pronounced among these gut microbiota-independent effects was an increase in the abundance of several digestive proteases in response to the egg white and brown rice proteins (Fig. 3). Interestingly, in a previous study focused on the fate of dietary protein in samples from Experiment 2, we found that dietary proteins from these two sources were recovered in large amounts in the stool due to inefficient digestion and absorption in the gut^14^. Taken together, these two findings indicate that dietary protein sources that are not efficiently processed in the gut induce changes in host digestive physiology, specifically by increasing the abundance of digestive proteases or host proteolytic activity independent of gut microbiota interactions. These proteases have been shown to have functions beyond digestion in the gut through their interactions with protease receptors and inhibitors, including extracellular matrix remodeling and maintaining gut immune homeostasis by performing proteolytic activity against chemokines and receptors^57^. Increased host protease activity in the gastrointestinal tract can lead to intestinal barrier disruption and mucosal damage and has been previously linked to inflammatory bowel diseases^58,59^.

The most pronounced microbiota-dependent effects were unsurprisingly immune-related, as germ-free mice are known to have underdeveloped immune systems^39,40^. Yeast protein led to an increase in the abundance of immune response proteins, several with antimicrobial functions, including IGHA, PIP, ZG16, and calprotectin subunit S100A9 (Fig. 4). Notably, increased abundance of calprotectin is used in clinical settings for IBD diagnosis and indicates onset of inflammation^60^. These microbiota-dependent immune responses in the yeast protein group were also associated with an increased abundance of overall bacterial proteins in the stool, reflective of a bacterial bloom reported previously^14^. However, interestingly, yeast protein was also associated with an increased abundance of REG3 lectins in both germ-free and conventional mice, and this specific antimicrobial innate immune response was not dependent on the gut microbiota. These results suggest that yeast protein, likely due to its microbial characteristics, induces the host’s innate immune response, an effect that is enhanced by either the interactions of yeast with the gut microbiota or the effects of the gut microbiota on the reactivity of the host’s immune response. Further investigating these yeast induced immune responses in the context of a complex diet is essential, as yeast and microbial protein found in fermented foods are regularly consumed in large amounts in human populations^33^.

We further found that host proteins involved in the maintenance of the intestinal epithelium and the gut barrier responded in both the presence and absence of the gut microbiota (Fig. 5). However, these proteins’ responses differed in magnitude or direction depending on the presence of the gut microbiota, indicating that, at least in part, host protein responses to diet depended on host-microbiota or diet-microbiota interactions (e.g.CEACAM1, CDHR5, MUC17 and TFF3). Most of these proteins responded strongly to egg white and yeast protein in both the germ-free and conventional mice.

Additionally, our results show that mice fed egg white protein had a significantly thinner mucus barrier, which was associated with a lower abundance of host mucus proteins and a higher abundance of microbial enzymes associated with mucin degradation (Fig. 6). These microbial effects were further associated with a higher abundance of *A. muciniphila* in mice fed egg white protein (Fig. 1G). Since microbial enzymes associated with mucin degradation can potentially degrade the mucus barrier, we hypothesized that these diet-microbiota interactions mediated egg white’s effect on the host mucus barrier. However, germ-free mice fed egg white protein also had a lower abundance of host mucus proteins than those fed brown rice protein, though the magnitude of this response was smaller than that in conventional mice, suggesting that some of the host mucus response to egg white protein might be microbiota-independent (Supp. Fig. 2). Therefore, we propose that egg white protein impacts the gut barrier via two separate mechanisms, one of which is dependent on the microbial-mediated degradation of the mucus layer, while the other is dependent on the impact of the increased host proteolytic activity on the gut barrier and epithelium independent of the microbiota.

Our results from the crossover feeding trial in humans showed that, despite high interindividual variability (Fig. 7B), the effects of dietary protein source on gut physiology and the microbiota seen in mice extend to humans consuming a far more complex diet than the defined diets fed to lab mice. We found that in humans, egg white protein was associated with an increased abundance of specific digestive proteases and gut epithelial proteins (Fig. 7E and F). In contrast, pea protein was associated with an increased abundance of antimicrobial proteins. Digestive proteases higher in the egg white protein group included pancreatic chymotrypsin-like elastase family member proteases, which were also higher in mice fed egg white protein. Interestingly, this increased proteolytic host response correlates with the reduced digestive efficiency of egg white proteins that we previously observed both in the same human samples^55^ and mice^14^. This suggests that harder to digest protein sources trigger a greater host proteolytic response in both mice and humans. Several gut epithelial maintenance proteins involved with cellular repair, stress response, apoptosis, and cellular adhesion, including annexin A5 and A7, programmed cell death 6 interacting protein, and claudin-7, were higher in the egg white protein group. Increased abundance of these proteins in stool could therefore reflect changes in epithelial turnover and barrier integrity in response to egg white protein. This interpretation is supported by the fact that subjects eating this protein source exhibited increased gut permeability^55^. On the other hand, pea protein consumption was associated with a higher abundance of carbonic anhydrase 4 (CA4) and secretory IgA related proteins (IGHA1, IGHA2 and JCHAIN) as well as PIGR, which transports IgA across the epithelium for secretion into the gut lumen^61^. These proteins help maintain a healthy gut barrier: CA4 by bolstering the mucus layer via bicarbonate production and IgA by defending the gut barrier from microbial invasion^62,63^.

In terms of gut microbial responses to egg white and pea protein in humans, while there were no differences in overall microbial taxonomic composition between the two groups, microbial function differed significantly (Fig. 7C and D). A previous analysis of this human data found that amino acid metabolism shifted significantly^55^. Our re-analysis revealed that the abundance of microbial enzymes associated with mucin degradation was significantly higher post egg white protein consumption compared to post pea protein consumption (Fig. 7H). Notably, we observed that this increase in the abundance of microbial enzymes with mucin-degrading potential in response to egg white protein was consistent across three independent datasets, including the two mouse experiments^13^ and this human study. However, unlike in the mice, the host mucus proteins did not differ between the egg white and pea protein groups in human stool samples, even as other epithelial and barrier proteins changed in abundance and intestinal permeability increased in the egg white protein group (Fig. 7G). These findings demonstrate that egg white protein consistently increases microbial mucin degradation potential across mice and humans, while its effects on the host mucus barrier manifest via variable readouts across species.

In summary, our results show that dietary protein source impacts host gut physiology beyond responses to nutrient content and that some of the effects are microbiota-dependent, while others are microbiota-independent. These findings provide key insights into how dietary protein sources mediate the differential health outcomes in response to dietary protein sources that have been observed in large-scale population studies^4,5^. Additionally, our study establishes a framework for identifying and characterizing site-specific protein-level gene expression responses in the gut using non-invasive stool proteomics. While our study captured host responses in stool, which mainly captures secreted and luminal host proteins, future work should investigate shifts in intracellular host proteins in intestinal epithelial cells. This would include proteins that impact cellular metabolism and respond to differences in nutrient and amino acid availability associated with different protein sources. Characterizing these responses could reveal intracellular processes driving changes in the secreted proteins measured in stool. Future studies should further investigate systemic host responses to dietary protein source and determine how shifts in other macronutrients such as fiber and fat change these host and microbial responses to specific dietary protein sources. With this study, we conclude our three-part series showing how metaproteomics is a powerful tool for analyzing dietary effects on the host and gut microbiome^13,14^. Previously, we showed that distinct protein sources have distinct fates in the intestine^14^, which are linked to changes in gut microbiome function^13^. Here we show that these protein sources also change host physiology. This provides a powerful framework for future research investigating how different macronutrient sources interact to affect microbiota function and host physiology, which can ultimately inform nutrition guidelines and therapeutic interventions for a diversity of diseases related to nutrition and the gut microbiota.

## Methods

### Animals and housing

In Experiment 1 (Fig. 1A), C57BL/6J mice (40 females, 4-5 weeks old, housed 4 mice per cage) purchased from Jackson Laboratories were used in the defined-diet feeding experiment. In Experiment 2 (Fig. 2A), previously unpublished host proteomics data generated alongside the dietary protein data in Awan et al. (2025) was used. Briefly, 12 germ-free (6 male, 6 female, NC State University gnotobiotic core) and 12 conventionally raised (6 male, 6 female, Jackson Labs, Bar Harbor) C57BL/J6 mice, aged 3-6 months, were used for the defined diet feeding study. The germ-free mice were housed in gnotobiotic isolators and were monitored for sterility. The male and female mice were housed separately in groups of three.

All animals were kept at a 12-hour dark/light cycle at an average temperature of 21°C and 35% humidity. The bedding was autoclaved, and sterile water (Gibco) was used for both germ-free and conventional mice. All diets were gamma-irradiated and vacuum packaged, and the same diet batches were used for germ-free and conventionally raised mice in Experiment 2. For use in gnotobiotic isolators, the outside of the vacuum-sealed diet packages was further sterilized using peracetic acid for 30 minutes before introduction into isolators to avoid autoclaving, which would have introduced differences between diets fed to germ-free and conventional mice. The animals were fed, weighed, and assessed daily by trained animal handlers, and all cage changes were performed in a laminar flow hood for conventional mice. All animal experiments were conducted in the Association for the Assessment and Accreditation of Laboratory Animal Care International (AAALAC) accredited Laboratory Animal Facilities at the NC State University College of Veterinary Medicine. The animal care protocol was approved by NC State’s Institutional Animal Care and Use Committee (Protocol # 18-034-B for conventionally raised mice and # 18-165-B for germ-free mice).

### Animal diets and sample collection

All defined diets used in this study were obtained from Inotiv (previously Envigo Teklad). Dietary components other than the source of the dietary protein did not change and were consistent and defined across the different diets. Diets were sterilized by gamma irradiation, vacuum-packaged, and sterilized on the outside before being introduced into the gnotobiotic isolators. The defined diets used in the experiment were isocaloric and contained all component macronutrients in the same proportions. The only difference between the diets was the source of the dietary protein. The diets contained purified protein isolates from a single source and were not supplemented with amino acids. The exact composition of the diets is available in Blakeley-Ruiz et al., 2025^13^. Mice were fed standard chow before starting the experimental diets.

In Experiment 1, four days before the start of the experimental diet regimen, a cage change and bedding swap were performed to ensure the microbiota of the mice between cages would be similar at the start of the experiment. To accomplish this, a small amount of soiled bedding was taken from each cage of mice, mixed in a sterile flask, and redistributed to each clean cage. On day 0, all mice were switched to experimental diets containing purified protein isolates from either 20% soy protein, 20% yellow pea protein, 20% torula yeast protein, 20% brown rice protein, or 20% egg white protein, with n=8 mice per diet group in 2 cages. The mice were fed the defined diets for 7 days, and stool pellets were collected on days 0 and 7. Animals were humanely euthanized by CO_2_ asphyxiation followed by cervical dislocation prior to necropsy. During necropsy, tissue snips were taken of the ileum, cecum, and colon and flash frozen in liquid nitrogen, then stored at −80°C until further analysis.

In Experiment 2, diets containing 20% soy protein, 20% casein, 20% brown rice protein, 40% soy protein, 20% torula yeast protein, 40% casein, 20% yellow pea protein, and 20% egg white were fed to each conventional and germ-free mouse *ad libitum* for 7 days per diet in this sequence. At the end of the dietary sequence, mice were divided into two groups, and one group was fed the 20% casein diet, and the other was fed the 20% soy protein diet again as a control to see if the effects observed at the beginning of the feeding sequence were reproducible. After feeding a defined diet for a week, stool samples were collected from each mouse before switching to the next diet. NAP preservation solution (935 ml autoclaved Milli-Q water, 700 g Ammonium Sulfate, 25 ml of 1 M Sodium Citrate, and 40 ml of 0.5 M EDTA adjusted to pH 5.2 using 1 M H_2_SO_4_) ^64^ was used to preserve and store the stool samples at a 1:10 sample-to-solution ratio. This method prevented sample modification, for example by proteases, and allows for rapid preservation of samples while working in gnotobiotic isolators in which liquid nitrogen cannot be used. Samples from germ-free mice and conventionally raised mice were treated identically to maintain consistency. Samples were roughly homogenized using a sterilized disposable pestle and then stored frozen at −80°C.

### Mucus Thickness Quantification

Colonic mucus thickness was quantified as previously described^65^. Briefly, mouse colons were harvested in Experiment 1, and colonic regions containing a stool pellet were fixed with fresh Carnoy’s fixative solution (methanol:chloroform:glacial acetic acid; 60:30:10) for 4 days, then sequentially washed in methanol, ethanol, and xylenes. Tissues were embedded in paraffin, sectioned (5 μm), and stained with Alcian Blue. Slides were digitized at 20X magnification using an Olympus VS-120 slide imager, and the mucus layer thickness was measured at ∼50 μm intervals along the circumference of each colon cross section (30 to 300 measurements per mouse) using NIH ImageJ in a blinded manner^66^. Per mouse, measurements from at least n=3 sections were averaged.

### Human study sample collection

Human stool samples from the randomized crossover human feeding trial in Experiment 3 (Fig. 7A) were collected in accordance with the protocols and procedures described in Thaker et al., 2026^55^. Briefly, healthy humans were randomly assigned to a group to eat one of two diets in which 70% of their daily protein was derived from either pea or egg white protein isolate for 10-12 days, followed by a 14-day wash-out period after which the participants crossed over to a different diet containing the alternative protein source for an additional 10-12 days. Stool samples from each individual were collected before starting the experimental diet and at the end of each feeding period. Stool samples were homogenized by adding 0.5-1 mL of UHPLC water and mixing with a hand-held tissue homogenizer (Omni Micro Homogenizer, 115 V, Omni International), then aliquoted and stored at −80°C until further metaproteomic analysis as previously described^55^. The study was approved by the Institutional Review Board at Stony Brook University (SBU) (IRB# 2022-00427) and registered on Clinicaltrials.gov (NCT05619939).

### Protein extraction from stool samples and peptide preparation

To extract proteins from stool and tissue samples from the mouse experiments, we first lysed the samples by bead-beating in 2 mL Lysing Matrix E tubes (MP Biomedicals) for 5 cycles of 45 s at 6.45 m/s with 1 min between cycles in SDT lysis buffer (4% (w/v) SDS, 100 mM Tris-HCl pH 7.6, 0.1 M DTT). Samples were heated to 95°C for 10 minutes, and the resulting lysates were centrifuged (21,000 x g, 5 min) to remove the lysing beads and any other debris.

To prepare peptides from lysates of all mouse and human samples, we followed the filter-aided sample preparation (FASP) protocol with modifications^67^. 60 µl of the lysate was mixed with 400 µL of UA solution (8 M urea in 0.1 M Tris/HCl pH 8.5). The mixture was loaded onto a 10 kDa 500 µL filter unit (Microcon Biomax PES 10 kDa filters) and centrifuged at 14,000 x g for 30 minutes. This step was repeated up to a total of 3 times to load the filter to capacity. The filters were washed using 200 µl of UA solution, centrifuged at 14,000 g for 40 min, and incubated with 100 µl IAA (0.05 M iodoacetamide in UA solution) for 20 min, followed by centrifugation at 14,000 *g* for 20 min. Filters were then washed with 100 uL of UA buffer 3 times. At this stage, the protocol was modified for Experiment 1 mouse and Experiment 3 human samples, and filters containing mouse tissue and stool samples were washed with 200 μL of ABC buffer (50 mM Ammonium Bicarbonate), followed by two washes with 200 μL of a 60% acetonitrile: water solution. All samples from all experiments were then washed with 100 µl of ABC (50 mM Ammonium Bicarbonate) buffer 3 times before adding MS-grade trypsin (Thermo Scientific Pierce, Rockford, IL, USA) solubilized in 40 µl of ABC buffer to the filters and incubating in a wet chamber at 37 °C to digest the proteins into peptides. 1 ug of trypsin was added to all the samples. After overnight digestion, peptides were eluted off the filter by centrifugation at 14,000 x g for 20 minutes. 50 uL of 0.5 M NaCl with 0.4% formic acid was added to the filters, followed by centrifugation at 14,000 x g for 20 minutes. The concentration of the resulting peptides was determined using the Pierce Micro BCA assay (Thermo Scientific Pierce) following the manufacturer’s instructions.

### LC-MS/MS analysis of peptides

Peptides from mouse stool and tissue samples from Experiment 1, and from human stool samples in Experiment 3, were analyzed in a block-randomized manner using 1D-LC-MS/MS on a Vanquish Neo UHPLC system (Thermo Fisher Scientific) coupled to an Orbitrap Astral (Thermo Scientific). 500 ng of peptide was loaded onto a PepMap™ Neo Trap Cartridge (5 μm particles, 300 μm X 5 mm, Thermo Fisher Scientific) and separated using an EASY-Spray™ PepMap™ RSLC C18 column (2 µm particles, 75 cm x 75 µm ID, Thermo Scientific) with the following 70 min gradient at a flow rate of 0.25 ul/min: 1%-5% B in 0.1 min then 5%-28% B in 49 min, 28%-40% B in 9 min, 40%-99% B in 2 min, then isocratic at 99% B for 10 min). The eluted peptides were ionized using an Easy Spray electrospray source (static spray voltage 2,000 V (+)) and analyzed using the Orbitrap Astral mass spectrometer (Thermo Fisher Scientific). Mass spectra were acquired in positive ionization mode using data-independent acquisition (DIA) with the following parameters: full MS1 scans in the Orbitrap with 380-980 m/z range, 240,000 resolution, 500% AGC target (equivalent to 2.5e7 ions), maximum injection time 5 ms, 1 microscan, profile mode, m/z 445.12003 lock mass. MS/MS scans were performed in the Astral analyzer with 150-2000 m/z range, 500% AGC target (equivalent to 2.5e5 ions), maximum injection time 5 ms with 1 microscans, isolation window 1 m/z, resulting in 600 scan events across the 380 to 980 m/z range, normalized collision energy of 32%, loop time of 1.5 s, and profile mode.

Peptides from Experiment 2 samples were analyzed using 1D-LC-MS/MS, as previously described in Blakely-Ruiz et al., 2025^13^. 600 ng of peptides from each of the mouse stool samples was loaded onto a 5 mm, 300 μm ID C18 Acclaim® PepMap100 pre-column (Thermo Fisher Scientific) with loading solvent A (2% acetonitrile, 0.05% TFA) in an UltiMate™ 3000 RSLCnano Liquid Chromatograph (Thermo Fisher Scientific). Peptides were separated on an EASY-Spray analytical column heated to 60°C (PepMap RSLC C18, 2 µm material, 75 cm× 75 µm, Thermo Fisher Scientific) using a 140 min gradient at a flow rate of 300 nl/min. The first 102 minutes of the gradient went from 95% eluent A (0.1% formic acid) to 31% eluent B (0.1% formic acid, 80% acetonitrile), then 18 min from 31 to 50% B, and the last 20 min at 99% B. A 100% acetonitrile wash run was performed between each sample to minimize carryover. The eluted peptides were ionized using an Easy-Spray source and analyzed in a Q Exactive HF hybrid quadrupole-Orbitrap mass spectrometer (Thermo Fisher Scientific) with the following parameters: m/z 445.12003 lock mass, normalized collision energy equal to 24, 25 s dynamic exclusion, and exclusion of ions of +1 charge state. Full MS scans were acquired for 380 to 1600 m/z at a resolution of 60,000 and a max IT time of 200 ms, and data-dependent MS^2^ was performed for the 15 most abundant ions at a resolution of 15,000 and max IT of 100 ms.

### Database construction, protein identification and quantification

The protein sequence database used to identify proteins in both mouse experiments (Experiments 1 and 2) consisted of dietary, host, and microbial proteins, as previously described in Blakeley-Ruiz et al., 2025^13^. Custom databases specific to each diet/sample type were used in line with previous recommendations^68^. For the host component of the database, the reference proteome for the C57BL/6J mouse (*Mus musculus*, UP000000589) was downloaded from UniProt and added to the dietary and microbial components.

The protein database for human stool samples was constructed using human, microbial, and dietary sequences as described previously in Thaker et al., 2026^55^. However, the microbial sequences were generated from a re-analysis of the 34 sample-matched metagenomes produced in the previous study. Briefly, metagenomes were re-assembled to produce MAGs. For the human microbiome data, we used the raw metagenomic reads from the 34 metagenomes sequenced in the previous publication^55^ to generate MAGs and protein sequences from these MAGs. For this, we trimmed reads with Trim_galore v 06.10 and removed host-read contaminants with bbmap v 39.01. We assembled reads by sample using MEGAHIT v 1.2.9^69^ with the following parameters: –k-max 141, –k-step 10, –min-contig-len 1000. Resulting contigs were combined into a single concatenated list of contigs and binned using SemiBin2 v 2.2.0^70^ using the SemiBin2 multi_easy_bin workflow, which included first mapping reads using bowtie v 2.5.4^71^. Bin quality was evaluated using CheckM2^72^, and we used dRep v3.4.5^73^ to cluster medium-quality bins into species groups using 95% ANI as the species cutoff^74,75^. We assigned taxonomy to these MAGs using GTDB-Tk v2.3.2 with the GTDB reference r214^76^. We predicted protein sequences from the concatenated assembly using Prodigal^77^ and clustered them at 95% using CD-HIT^78^. Protein groups that could be linked to a MAG were assigned to that MAG’s species group if all the MAGs in the protein group belonged to the same species. The dietary sequences were compiled from the proteomes and protein sequences of 242 common food species^79^. Every set of protein sequences (human, microbiota, individual dietary components) was independently clustered with a 95% identity threshold using cd-hit. The final database contained 49,042 human, 1,311,254 microbial, and 3,759,297 dietary protein sequences.

The MS^2^ spectra from the DIA-analyzed Experiment 1 and Experiment 3 samples were searched using DIA-NN 2.2^80^ with a predicted spectral library generated *in silico* using the respective sample-specific databases described above and the following parameters: trypsin digest, maximum 1 missed cleavage, and carbamidomethylation as a fixed modification. Protein inference and grouping were performed at the isoform level, and precursors and proteins were filtered at an FDR of 5%. The cross-run normalized MaxLFQ quantification values in the protein group matrix output file were used for all downstream analyses.

The MS^2^ spectra from the DDA analyzed Experiment 2 samples were searched using the sample-specific database for each diet group using run calibration and the Sequest HT node in Proteome Discoverer software version 2.3 (Thermo Fisher Scientific) with the following settings: trypsin (Full), maximum 2 missed cleavages, 10 ppm precursor mass tolerance, 0.1 Da fragment mass tolerance, and maximum 3 equal dynamic modifications per peptide. Dynamic modifications, including oxidation on M (+15.995 Da), deamidation on N, Q, R (0.984 Da), and acetylation on the protein N-terminus (+42.011 Da), and the static modification carbamidomethyl on C (+57.021 Da) were used. Proteins were quantified based on the area under the curve method in the Minora Feature Detector node with the following parameters: 5 minimum number of non-zero points in a chromatographic trace, 0.2 min maximum retention time of isotope pattern multiplets, and high PSM confidence^81^. The Percolator node in Proteome Discoverer was used to calculate peptide false discovery rate (FDR) with the following parameters: maximum Delta Cn 0.05, a strict target FDR of 0.01, a relaxed target FDR of 0.05, and validation based on q-value. The precursor ion quantifier node was used for quantification using the following settings: unique and razor peptides used for quantification and precursor abundance based on area. Only master proteins using the 5% FDR cutoff were included in the downstream data analysis.

### Host proteome data analysis and visualization

Host proteins were extracted from the stool and tissue metaproteomes, and only host proteins present in at least 70% of all the replicates of one diet condition were retained for normalization. Normalization was done by dividing the abundance of each host protein that passed the prevalence filter by the summed abundance of all host proteins in that sample, to obtain the relative abundance of each host protein as a percentage of the total host proteome i.e, organism-specific total sum scaling, or orgTSS%.The resulting protein abundances were transformed using a centered log-ratio transform in the R package compositions (version 2.0.5)^82^ and used for the principal coordinate analysis and the PERMANOVA in Fig. 1B, Fig. 2B-D, and Fig. 7B.

For the concordance analysis in Fig. 1, we assessed whether the host responses in the tissues are captured in the stool as well. To do this, we compared the 5 experimental diets in Experiment 1 in a pairwise manner across 10 total pairwise comparisons for each of the 3 tissue and stool proteomes. For the pairwise diet comparisons, we first filtered out proteins that were not present in at least 50% of replicates in both groups. The abundances of the remaining proteins were normalized using total sum scaling. Then, for each protein, using the log2 ratio of its mean abundance, we calculated the log2 fold change in its abundance between the two diets being compared. Since there were more host proteins detected in the tissues than in the stool, for the concordance analysis we only included proteins with a log2 fold change value in both the stool and the tissue being compared. Then, to determine whether the response of each protein to dietary perturbation in the ileum, cecum, and colon was captured in the stool irrespective of the specific diets being compared, we correlated its fold change in the stool to its fold changes in each of the 3 tissues across all dietary comparisons (Fig 1C). Only proteins that changed in abundance in both the stool and tissue in at least 6 dietary comparisons were included in this analysis because correlation analysis on fewer data points can lead to unreliable results.

We also quantified the level of directional agreement between the stool and tissue log2 fold change for each protein by calculating what percentage of dietary comparisons for that protein had the same direction of change in the stool and the tissue (% same sign, Fig 1C). To further assess how concordant the overall host response between tissue and stool was for each dietary comparison in the ileum, cecum, and colon, we correlated the log2 fold change in the abundance of all proteins in a dietary comparison in the stool and each of the three tissues (Fig 1D). To focus on host proteins responding to dietary perturbation, we only included proteins with a log2 fold change of at least 1 in both the stool and the tissue. ANOVA, followed by Tukey’s HSD post hoc test (rstatix package in R)^83^, was used to identify differentially abundant proteins across groups using stool host proteomics data from Experiment 2 for Figures 2-5.

For the human crossover feeding trial in Experiment 3, all pre- and post-diet intervention comparisons were made within each individual. For each participant, we calculated a log2 fold change (log2FC) for each protein between the post and pre timepoints within each diet treatment and between the post points between diets. We then averaged these individual fold changes to determine the within-diet response (Fig. 7E). We tested between-diet effects in two ways to identify proteins whose response differed between diets, both in terms of difference in change from baseline and at post intervention. First, we used a linear model of individual post vs pre log2FC on diet, testing whether the magnitude of the post vs pre change in each diet differed between diets. Then we used an ANCOVA of post-intervention abundance adjusted for pre-intervention abundance, with individual as a random intercept, testing whether post intervention abundance differed between diets after accounting for differences in baseline.

Based on the outcome of these two between-diet analyses, we classified proteins as diet responders if they had a nominally significant diet effect (p <0.05) in either the linear model of individual post vs pre log2FC on diet, or in the ANCOVA of post-intervention abundance adjusted for baseline. We then tested between-diet differences at the post timepoint for these diet responder proteins and for the most abundant stool host proteins which had a mean relative abundance of at least 1% in any diet and timepoint group, using a paired t-test with Benjamini-Hochberg corrected p-values (Fig. 7F). Protein functions were annotated and classified using UniProt, Panther, Gene Ontology, and an extensive literature review. R (R version 4.2.2)^84^ and Excel were used for all data processing and analysis. R (ggplot2, version 3.5.1)^85^ and Adobe Illustrator were used for data visualization.

### Microbial data analysis and visualization

For the mouse microbial data, proteins had already been assigned to species groups in a previous publication based on metagenome-assembled genomes (MAGs)^13^. For the human microbial data, we used the species group assignments generated in the re-analysis of the metagenomic data described in the database construction section. Proteomic species (MAG) abundance was calculated using a modified version of the species biomass approach, adapted for DIA data^86,87^. We only considered protein groups with a single isoform identifier, unique to a single species, and we summed the QuantUMS^88^ intensity of all the proteins belonging to a single species to calculate biomass. Biomass was normalized to a per-sample percentage made up of only the microbiome fraction of the proteins. To assess species composition, the Bray-Curtis dissimilarity index was calculated using the vegan R package^36^. We then assessed the community using the PCoA and PERMANOVA functionalities within the package and visualized them with ggplot2. For functional analysis, of both the human and mouse data. The identified proteins were assigned KEGG orthologous groups using MANTIS^89^. We summed the abundance of proteins by the KEGG orthologous group to assess shifts in overall microbiota function and performed the same compositional analyses as described for species groups.

## Supporting information

Supplemental Results

## Ethics statement

NC State’s Institutional Animal Care and Use Committee approved the animal care protocol (Protocol # 18-034-B for conventionally raised mice and # 18-165-B for germ-free mice). The human diet study was approved by the Institutional Review Board at Stony Brook University (SBU) (IRB# 2022-00427) and registered on Clinicaltrials.gov prior to recruitment of subjects (NCT05619939).

## Data availability

Proteomics data, including raw mass-spectrometry files and databases used, were deposited to the ProteomeXchange Consortium via the PRIDE partner repository with the dataset identifier PXD082298 for Experiment 1 (reviewers can access using the project accession: PXD082298 and the token: uDYNTvOgzmQj), PXD041586 for Experiment 2 (https://www.ebi.ac.uk/pride/archive/projects/PXD041586), and PXD082075 (reviewers can access using the project accession: PXD082075 and the token: Dm19jh6ZY9Gm) for the human feeding trial.

## Supplemental materials

**Supplemental Table 1:** Raw metaproteomic data for all host, microbial, and dietary proteins detected in all tissue and stool samples analyzed in Experiment 1.

**Supplemental Table 2:** Table showing total sum scale normalized (orgTSS%) abundance of host proteins identified in at least 70% of the replicates in all stool samples in Experiment 2.

**Supplemental Table 3:** Table showing the protein functional categories in Fig. 1E, determined using Panther protein class and Gene Ontology.

**Supplemental Table 4:** Table showing total sum scale normalized (orgTSS%) abundance of host proteins in all stool samples from the crossover human feeding trial in Experiment 3.

**Supplemental Table 5:** Table showing the raw abundance values of annotated microbial proteins in stool samples from the crossover human feeding trial in Experiment 3.

**Supplemental Table 6:** Table showing the biomass values of microbial taxa quantified in stool samples from the crossover human feeding trial in Experiment 3.

**Supplemental Table 7:** Table showing the metadata associated with the stool samples generated in the crossover human feeding trial in Experiment 3.

## Author contributions

**AA**: Conceptualization, Data curation, Formal analysis, Investigation, Methodology, Visualization, Writing – original draft, Writing – review & editing.

**JABR**: Conceptualization, Formal analysis, Investigation, Data curation, Visualization, Writing – original draft, Writing – review & editing.

**CG**: Investigation, Data curation, Writing – review & editing.

**ST**: Formal analysis, Data curation, Visualization, Writing – original draft, Writing – review & editing.

**HC**: Investigation, Data curation, Writing – review & editing.

**AB**: Investigation, Data curation, Writing – review & editing.

**LD:** Investigation, Data curation.

**JCM:** Investigation, Data curation.

**DCM**: Funding acquisition, Resources, Supervision, editing of manuscript.

**MK**: Conceptualization, Data curation, Formal analysis, Funding acquisition, Investigation, Methodology, Supervision, Writing – original draft, Writing – review & editing.

## Acknowledgments

The authors would like to thank Dr. Casey Theriot and Cypress Perkins for their guidance, time, and resources in conducting the mouse experiments. The authors would also like to thank Dr. Amanda Ziegler for several consultations regarding data analysis strategies. All LC-MS/MS measurements were made in the Molecular Education, Technology, and Research Innovation Center (METRIC) at North Carolina State University. The Gnotobiotic Core at the College of Veterinary Medicine, North Carolina State University, is supported by the National Institutes of Health funded Center for Gastrointestinal Biology and Disease, NIH-NIDDK P30 DK034987. This work was supported by the National Institutes of Health under Award Numbers R35GM138362, R01DK118024, and K01DK144672 and the Pulse Crop Health Initiative of the US. Department of Agriculture (58-3060-2-034). The content is solely the responsibility of the authors and does not necessarily represent the official views of the National Institutes of Health.

