## Supplemental Results for "Dietary protein source alters gut physiology through direct and microbiota-mediated effects"

### Supplementary Results

***Difference in sex or gut microbial composition leads to differential host fecal proteome in response to specific dietary protein sources***

To determine the overall differences in the host fecal proteome in germ-free and conventional male and female mice fed different sources of dietary protein, we used hierarchical clustering based on Euclidean distances and the ward.D2 clustering method (pheatmap package in R)[1]. We found that the yeast, brown rice, and egg white protein groups formed distinct clusters in the germ-free and conventional mice. In the conventional mice, the plant-based dietary protein sources (soy, pea, and brown rice) grouped within the same overall cluster, and the animal-based dietary protein sources (casein and egg white) clustered together, while the microbial/fungal yeast protein formed its own cluster. This pattern was not observed in germ-free mice, where the egg-white cluster was closer to pea and soy, and casein and brown rice formed a separate cluster. Separation between the male and female host fecal proteomes was observed in the conventional mice fed yeast, egg white, brown rice, and casein protein diets, but no separation between the male and female mice was observed in the germ-free group. No separation was observed between the 20% and 40% soy and casein protein groups in either the germ-free or the conventional mice.

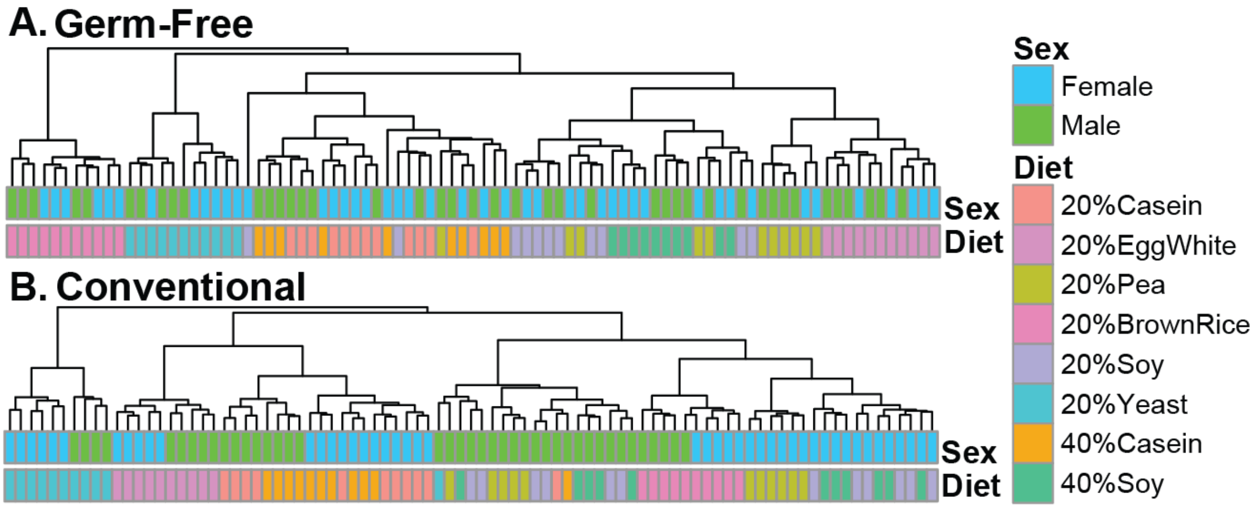


**Supplemental Figure 1. Dietary protein source differentially impacts host fecal proteome in germ-free and conventional mice.** Hierarchical clustering analysis of host fecal proteomes of germ-free and conventional, male and female mice fed defined diets containing 20% purified dietary protein from casein, egg white, pea, brown rice, soy, and yeast and 40% protein from casein and soy. Euclidean distances and the Ward.D2 clustering method from the pheatmap package in R were used to do the analysis.

***Dietary protein source impacts host proteins associated with the mucus layer in both the presence and absence of the gut microbiota***

We compared the summed relative abundance of stool host proteins associated with the mucus layer in germ-free and conventional mice fed purified dietary proteins from different plant and animal sources (Experiment 2, Fig. 2A) to determine the effect of dietary protein from different sources on the gut mucus barrier and the role of the gut microbiota in mediating these effects. We found that germ-free mice fed casein and brown rice protein had the highest summed abundance of host mucus proteins, while conventional mice fed egg white protein had the lowest abundance of these proteins.

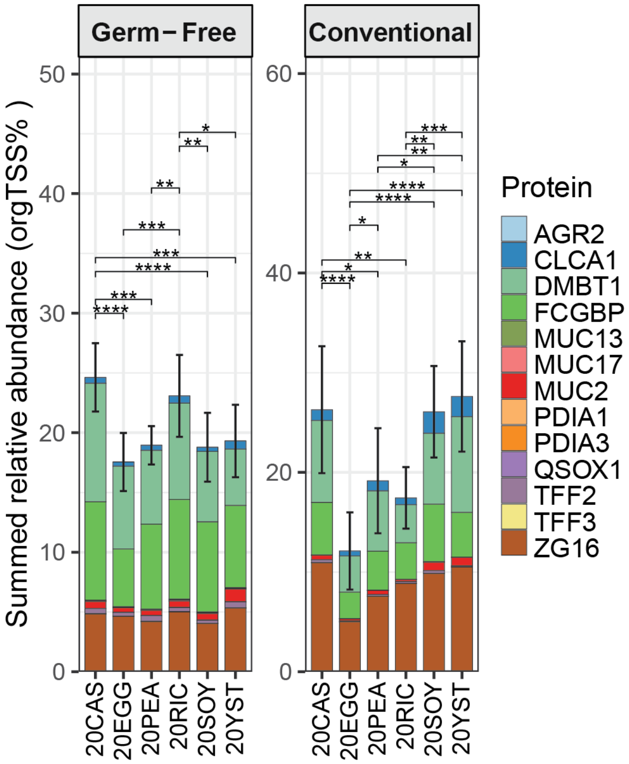


**Supplementary Figure 2**. **Dietary protein source impacts the abundance of host mucus-associated proteins in both germ-free and conventional mice**. Summed relative abundance of host proteins associated with the mucus layer in germ-free and conventional mice from Experiment 2. Each stack is the mean relative abundance of a protein, and the stacked bars represent the summed mean relative abundance of all mucus-related proteins in each diet. Statistical significance, determined using a t-test followed by Benjamini-Hochberg correction for multiple tests, is indicated by horizontal lines and asterisks (∗= p<0.05, ∗∗ = p<0.01, ∗∗∗ = p<0.001, ∗∗∗∗ = p<0.0001). Bars represent standard deviation from the group means.

***The abundance of host proteins associated with the mucus layer in stool does not differ between individuals fed pea versus egg white protein***

We compared the abundance of host mucus proteins in the stool of individuals fed diets supplemented with pea or egg white protein pre- and post-dietary intervention and found no difference at the level of individual proteins or in their summed relative abundance between the two diets post-intervention. However, we found that ZG16 increased significantly as a result of egg white protein from pre to post intervention ($p=0.05$). While not significant, we did see an increase in the abundance of CLCA1, QSOX1 and MUC13 from pre to post intervention in the pea group.

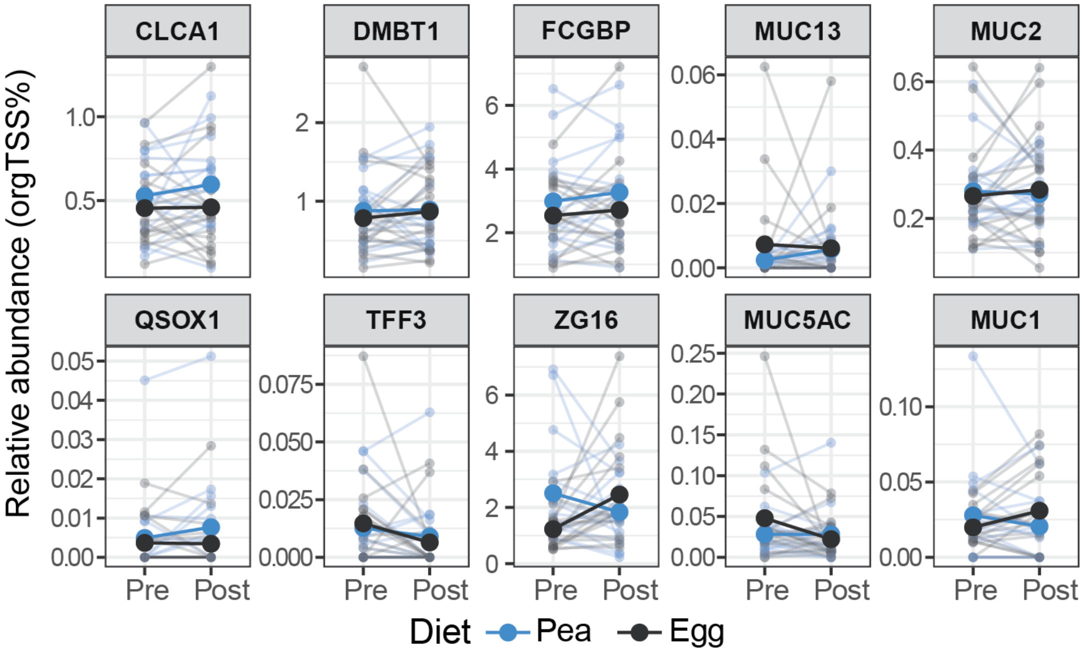


**Supplementary Figure 3. Host mucus proteins do not change significantly in response to egg white or pea protein supplementation in humans.** Relative abundance (orgTSS%) of host proteins associated with the mucus layer in stool from humans before and after consuming diets supplemented with pea or egg white protein in a randomized crossover feeding trial (Fig. 7A). Faint lines connect pre and post intervention measurements from one individual, and the solid bold points and lines represent the mean of each diet group. None of the individual proteins were significantly different between the two dietary groups post intervention. However, egg white protein led to a significant increase in the abundance of ZG16 from pre to post intervention.
